# Translation fidelity and respiration deficits in CLPP-deficient tissues: Mechanistic insights from mitochondrial complexome

**DOI:** 10.1101/2023.09.29.560101

**Authors:** Jana Key, Suzana Gispert, Gabriele Koepf, Julia Steinhoff-Wagner, Marina Reichlmeir, Georg Auburger

## Abstract

Mitochondrial matrix peptidase CLPP is crucial during cell stress. Its loss causes Perrault syndrome type 3 (PRLTS3) with infertility, neurodegeneration and growth deficit. Its target proteins are disaggregated by CLPX, which also regulates heme biosynthesis via unfolding ALAS enzyme, providing access of pyridoxal-5’-phosphate (PLP). Despite efforts in diverse organisms with multiple techniques, CLPXP substrates remain controversial. Here, avoiding recombinant overexpression, we employed complexomics in mitochondria from three mouse tissues to identify endogenous targets. CLPP absence caused accumulation and dispersion of CLPX-VWA8 as AAA+ unfoldases, and of PLPBP. Similar changes and CLPX-VWA8 comigration were evident for mitoribosomal central protuberance clusters, translation factors like GFM1-HARS2, RNA granule components LRPPRC-SLIRP, and enzymes OAT-ALDH18A1. Mitochondrially translated proteins in testis showed reductions to <30% for MTCO1-3, misassembly of complex-IV supercomplex, and accumulated metal-binding assembly factors COX15-SFXN4. Indeed, heavy metal levels were increased for iron, molybdenum, cobalt and manganese. RT-qPCR showed compensatory downregulation only for *Clpx* mRNA, most accumulated proteins appeared transcriptionally upregulated. Immunoblots validated VWA8, MRPL38, MRPL18, GFM1 and OAT accumulation. Coimmunoprecipitation confirmed CLPX binding to MRPL38, GFM1 and OAT, so excess CLPX and PLP may affect their activity. Our data elucidate mechanistically the mitochondrial translation fidelity deficits, which underlie progressive hearing impairment in PRLTS3.

## 1. Introduction

LONP1 and CLPXP are the two soluble proteases of the mammalian mitochondrial matrix, with extreme conservation until bacteria and plant chloroplasts [1, 2]. LONP1 is essential for life [3], plays the crucial role for protein turnover [4], combines the AAA+ unfoldase and the peptidase domain within one protein [4, 5], and assembles into homohexameric rings upon overexpression, purification and electron microscopy analysis, thus achieving maximal activity [6]. In comparison, CLPP absence may even extend lifespan in eukaryotes [7]; its functions become necessary only in periods of cell stress [1], when it acts in response to stalled ribosomes and possibly in the unfolded protein response of mitochondria (UPRmt) [8, 9]. CLPP by itself exerts only a chymotrypsin-like activity towards small peptides and depends on interaction with the AAA+ unfoldase / disaggregase CLPX for substrate recognition and unfolding, as well as activation of its proteolytic capacity [10, 11]. Bacterial ClpX was shown to perform chaperone roles independent from ClpP, cooperating with the Hsp70 family of chaperones (DnaJ/DnaK/GroEL) in the HssR defense system against heme-metal toxicity [11–15].

Mutations in human CLPP cause Perrault syndrome type 3 (PRLTS3), an autosomal recessive disorder characterized by complete infertility due to female primary ovarian insufficiency and male azoospermia after meiotic arrest at diplotene, with subsequent sensorineural deafness, followed by insidious neurodegeneration manifesting as ataxia, leukodystrophy and neuropathy [16, 17]. Most variants of Perrault syndrome are triggered by the failure of mitochondrial RNA processing and translation, as exemplified by other causal mutations in the tRNA-amino acid synthases HARS2 and LARS2, in the mitoribosomal factor RMND1, in the RNA degradation factor PRORP, in the nucleoid replication/repair factor TWNK, or in CLPP [18, 19].

In contrast, mutations in human CLPX trigger hematological disorders due to errors in heme biosynthesis [20, 21]. In zebrafish, the knockdown of CLPX can be rescued by supplementation of the heme precursor ALA (delta-amino levulinic acid) [22]. A common iron/heme-related pathway connects the two diverse clinical manifestations, given that porphyrias can progress into ataxia / leukodystrophy [23–29]. The relevant CLPX functions independent from CLPP were uncovered in *Saccharomyces cerevisiae* (where CLPP is not conserved) by a genetic screen, which showed that the CLPX ortholog activates the heme biosynthesis enzyme ALAS (5-aminolevulinic acid synthase), so CLPX deletion causes a 5-fold reduction of ALA and a 2-fold reduction of heme [30]. This study also reported that CLPX achieves this activation via controlling the incorporation of its cofactor, pyridoxal-5’-phosphate (PLP) into ALAS. PLP is a vitamin B6 derivative, which mediates Schiff base formation after a nucleophilic attack by lysine within the enzyme active site, but also has a chaperone role in protein folding [31]. This CLPX role for PLP access to ALAS and folding control of this enzyme is conserved until mammals [30]. Cells modulate the toxicity of excess porphyrin intermediates by very tight regulation of ALAS expression, in dependence on iron availability. The completion of heme synthesis is governed further downstream again in mitochondria via CLPX modification of protoporphyrin IX oxidase (PPOX) activity and ferrochelatase (FECH) levels. For chlorophyll biosynthesis, plant cells also have feedback from ALA levels onto the crucial porphyrin activation by magnesium chelatase [32, 33]. Plant magnesium chelatase protein sequence shows a coexistence of AAA+ and VWA domains, which appeared first in *Synechocystis* cyanobacteria within the catalytic ChlD subunit, and is still conserved in the mitochondrial VWA8 unfoldase of mammals despite their lack of chlorophyll production [34].

Bacterial ClpP can interact also with other AAA+ unfoldases from the Hsp100 family such as ClpA [35, 36], but in mammals, only the AAA+ unfoldase CLPB remains conserved and no stable interaction has been observed between ClpX and ClpB in bacteria [36]. AAA+ domains typically adapt a homohexamer conformation for maximal efficiency, whereas ClpP assembles into homoheptameric rings [37, 38], and a fully assembled ClpXP degradation machine in *E. coli* was shown to adopt a barrel-like form: two ClpP rings in the center represent the proteolytic chamber, while ClpX rings on either side are responsible for the targeting and linearization of unfolded or aggregated substrates [39]. The central cavity within the hexameric AAA+ ring can provide a tunnel through which RNA or DNA can be threat to alter the assembly or remove associated proteins [1, 34]. As first question of this study, we investigate whether mammalian AAA+ unfoldases other than CLPX interact physically or functionally with CLPP within the same folding pathways? Perhaps another AAA+ unfoldase assumes the workload when the CLPXP apparatus is dysfunctional. There are several candidates with known roles for mitochondrial protein complex dynamics and degradation in other contexts. They include SPATA5 and presumably VWA8 (given its BIOGRID interaction profile) in mitoribosomal and respiratory chain remodeling [40], BCS1L in respiratory complex III assembly [41], YME1L1, AFG3L2 and SPG7 in the mitochondrial inner membrane [42], as well as VCP and ATAD3 exerting unfolded protein response (UPR) tasks in the membranes of endoplasmic reticulum and mitochondria [43, 44].

As main targets of the CLPXP degradation machinery in bacteria and mammalian mitochondria during cell stress, incomplete translation products at stalled mitoribosomes are thought to be disaggregated and degraded. It was demonstrated that CLPP deficiency in mice causes impaired translation fidelity at mitoribosomes, claiming that this occurs due to the accumulation and inappropriate interactions of the rRNA chaperone ERAL1, which acts mainly in the assembly of mtSSU, the small mitoribosomal subunit [45–47]. However, the crucial role of ERAL1 for mitoribosome pathology in CLPP-null mice has since been disputed [48, 49]. Problems of translation fidelity in mitoribosomes would affect primarily the 13 proteins encoded by mitochondrial DNA to function within respiratory chain complexes [50]. Indeed, it is clear that the respiratory complex-IV is prominently affected in CLPP-null mice. Although respiratory activity in the brain and muscle appeared almost normal, the complex-IV activity in the liver was halved, presumably due to reduced abundance of mitochondrially encoded MTCO1 protein (30% in prominently affected testis), despite a *mtCo1* (*Cox1*) mRNA increase in all tissues studied (5-fold in testis) [51]. The biosynthesis of MTCO1 protein is preferentially prone to mitoribosomal stalling, given that its sequence contains 1 triproline and 3 diproline motifs in humans via rodents to zebrafish, 3 diprolines in *Drosophila melanogaster* flies, and 2 diprolines in *Saccharomyces cerevisiae* yeast [52]. All other mitochondrially encoded proteins have maximally 2 diprolines in their sequence (MTCO3 and MT-ATP8), and reduced abundance in CLPP-null mice was only observed for MTCO1 and MTND6 in some tissues [51]. Beyond translation impairment, preliminary data also suggested a generalized reduction of respiratory supercomplexes in CLPP-null mouse heart and liver tissue [51]. Altered disassembly and decreased turnover was documented in detail for the respiratory complex-I peripheral arm in CLPP-null mice [53]. Thus, assembly/disassembly dynamics may be affected not only for the mitoribosome and the respiratory chain, but also for other protein/RNA complexes as a secondary consequence of CLPP deficiency.

Therefore, a second question of this study focuses on the identification of CLPXP folding/degradation targets, to understand the molecular mechanisms of Perrault syndrome. Previous efforts to define the prominent CLPXP degradation substrate proteins in the matrix of bacteria/chloroplasts/mitochondria have yielded widely variable results in different species, and there is an ongoing debate if metabolic enzymes [54, 55] or ribonucleoproteins [49] are preferentially targeted. Substrate trap assays with overexpressed mutant CLPP, as well as global proteome profiles in tissues and cells without stress or after stressor administration, and finally databases on the protein interactions of overexpressed or endogenous CLPX have been scrutinized [49], without succeeding to pinpoint an invariably CLPXP-dependent protein. Recent studies of CLPP-null mice and CLPP-mutant patient cells showed the unfoldase CLPX to co-accumulate with selected protein subunits of the mitoribosome and the RNA granule, such as the translation elongation GTPase GFM1, the mitochondrial mRNA processing factor LRPPRC (and at least in mouse also the G-quadruplex RNA unwinding factor GRSF1), as well as the CLPX stabilizer POLDIP2 [45, 56]. Together with CLPX, accumulation was shown also for the molecular chaperone HSPA9, and the amino acid biosynthesis enzyme OAT, so they were interpreted as CLPP substrate candidates [57, 58].

Avoiding overexpression of exogenous proteins and thus aberrant interactions, two-dimensional blue native gel electrophoresis (BNE) with mass spectrometry (MS) of consecutive gel slices (dubbed as “complexomics”) was employed in two pioneer studies to define the endogenous physiological interactors of CLP-related AAA+ unfoldases. A complexome study in *E. coli* failed to detect ClpX, but showed ClpB to comigrate with the molecular chaperone DnaK, the translation initiation factor InfB, and the small ribosomal subunit mRNA unfoldase RpsA [59]. A preliminary complexome profile of one CLPP-deficient mouse heart suggested accumulation and disperse migration of CLPX (6-fold) in parallel to the CLPX stabilizer POLDIP2, RNA granule factors GRSF1, TBRG4/FASTKD4 and LRPPRC, translation elongation GTPases GFM1 and GFM2, as well as the AAA+ unfoldase VWA8 [49, 53]. It remained unclear whether these observations can be reproduced in more samples and be extended to more severely affected tissues of CLPP-mutant mice or patients. Furthermore, any direct impact of CLPXP on mitoribosomes was not elucidated. While the complete 55S mitoribosome migrates at 2.7 MDa, the 39S mtLSU appears at 1.7 MDa, and the 28S mtSSU around 1 MDa. The analysis of additional CLPP-null mouse tissues regarding CLPX comigration and abnormal migration patterns of the endogenous proteins is useful to elucidate the primary events of physiological and non-physiological interactions as well as CLPX targeting selectivity.

**Figure 1.**
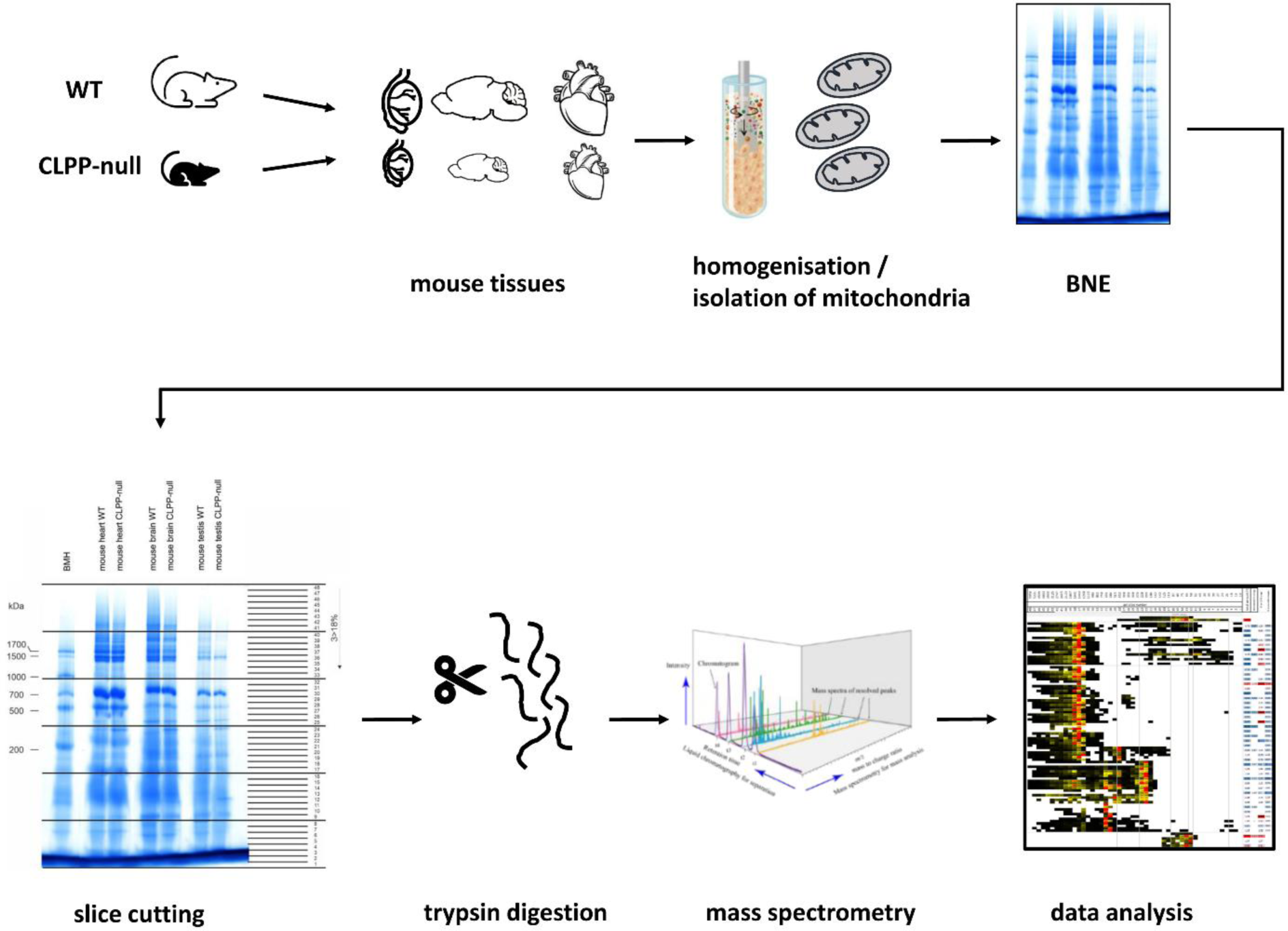
Scheme of experimental flow and data analysis, to elucidate CLPP-null triggered anomalies in CLPX interactions with endogenous protein complexes. Such complexome profiling efforts were now performed, using protein extraction with a low concentration of the mild detergent digitonin (so that respiratory supercomplexes and mitoribosomal subunits can remain assembled), followed by BNE and label-free mass spectrometry in 48 sections to quantify the abundance of proteins at each migration position (scheme of the experimental setup in Figure 1). We chose to study CLPP-null testis as maximally affected tissue with the broadest expression profile, the brain as modestly affected tissue with broad expression profile, and the heart muscle despite a lack of clinical affection and its limited expression profile for the sake of comparison with the previous complexome report. In the present manuscript, the focus was placed on AAA+ unfoldases and RNA processing factors to define the primary events of pathogenesis in PRLTS3. Secondary and compensatory events such as dysregulations of amino acid metabolism enzymes will be the subject of a subsequent manuscript.

## 2. Experimental procedures

### 2.1. Animal breeding, aging, and dissection

The generation of the CLPP-null (*Clpp*^−/−^) mice was described in detail before [51], and pups were bred from heterozygous matings. Mice were housed under specific-pathogen-free conditions under a 12 h light cycle with food and water *ad libitum* in the central animal facility (ZFE) of the University Hospital Frankfurt. The study was conducted according to the guidelines of the Declaration of Helsinki, and approved by the Institutional Review Board of Regierungspräsidium Darmstadt (protocol code V54-19c20/15-FK/1073, date of approval Sept. 28, 2016). Due to the complete infertility of CLPP-null homozygous mice of both sexes, breeding was performed simultaneously among multiple pairs of heterozygous mutation carriers, aging them under identical conditions until sacrificed for analysis. WT and CLPP-null mutant mice aged to the age of 8 weeks were sacrificed by cervical dislocation and used for organ dissection. Tissues were snap-frozen in liquid nitrogen and stored at -80 °C.

### 2.2. Experimental design and statistical rationale

For complexome investigations 1 WT as control vs 1 *Clpp*-null mouse were used. Mouse brain, heart, and testes tissues were taken for further processing. The results were not analyzed in a statistical manner, but the segregation of protein complexes, super-complexes and assembly intermediates was investigated and compared for consistency between different tissues.

### 2.3. Isolation of mitochondria in sucrose gradients

Mitochondria were resuspended in solubilization buffer (50 mM imidazole pH 7.50 mM NaCl, 1 mM EDTA, 2 mM aminocaproic acid) and solubilized with 20% (w/v) digitonin (Serva, Heidelberg, Germany). Samples were supplemented with 2.5 µl 5% Coomassie G250 in 500 mM aminocaproic acid and 5 µl 0.1% Ponceau-S in 50% glycerol. Equal protein amounts of samples were loaded on top of a 3 to 18% acrylamide gradient gel (dimension 14×14 cm). After native electrophoresis in a cold chamber, blue-native gels were fixed in 50% (v/v) methanol, 10% (v/v) acetic acid, 10 mM ammonium acetate for 30 min and stained with Coomassie (0.025% Serva Blue-G, 10% (v/v) acetic acid) [60].

### 2.4. Sample preparation for complexome profiling

Each lane of a BNE gel was cut into equal fractions and collected in 96 filter well plates (30-40 µm PP/PE, Pall Corporation). The gel pieces were destained in 60% methanol, 50 mM ammoniumbicarbonate (ABC). Solutions were removed by centrifugation for 2 min at 600×g. Proteins were reduced in 10 mM DTT, 50 mM ABC for one hour at 56 °C and alkylated for 45 min in 30 mM iodoacetamid. Samples were digested for 16 h with trypsin (sequencing grade, Promega, Madison, Wisconsin, USA) at 37 °C in 50 mM ABC, 0.01% Protease Max (Promega) and 1 mM CaCl_2_. Peptides were eluted in 30% acetonitrile and 3% formic acid, centrifuged into a fresh 96 well plate, dried in speed vac and resolved in 1% acetonitrile and 0.5% formic acid.

### 2.5. Setting HPLC / MSMS methods

Liquid chromatography / mass spectrometry (LC/MS) was performed on Thermo Scientific™ Q Exactive Plus equipped with an ultra-high performance liquid chromatography unit (Thermo Scientific Dionex Ultimate 3000) and a Nanospray Flex Ion-Source (Thermo Scientific, Waltham, MA, USA). Peptides were loaded on a C18 reversed-phase precolumn (Thermo Scientific) followed by separation on a 2.4 µm Reprosil C18 resin (Dr. Maisch GmbH, Ammerbuch, Germany) in-house packed picotip emitter tip (diameter 100 µm, 15 cm from New Objectives, Littleton, MA, USA) using a gradient from 4% ACN, 0.1% formic acid to 25% eluent-B (99% acetonitrile, 0.1% formic acid) for 30 min followed by a second gradient to 50% B with a flow rate 300 nl/min and washout with 99% B for 5 min. MS data were recorded by data-dependent acquisition. The full MS scan range was 300 to 2000 m/z with a resolution of 70000, and an automatic gain control (AGC) value of 3E6 total ion counts with a maximal ion injection time of 160 ms. Only higher charged ions (2+) were selected for MS/MS scans with a resolution of 17500, an isolation window of 2 m/z and an automatic gain control value set to E5 ions with a maximal ion injection time of 150 ms. MS1 Data were acquired in profile mode.

### 2.6. Data analysis with MaxQuant

MS Data were analyzed by MaxQuant (v1.6.17.0) [61] using default settings. The search engine was integrated in MaxQuant. Proteins were identified using mouse reference proteome database UniProtKB with 55470 entries, released in 4/2021. The enzyme specificity was set to Trypsin with a maximal number of missed cleavages of 2. Acetylation (+42.01) at N-terminus and oxidation of methionine (+15.99) were selected as variable modifications and carbamidomethylation (+57.02) as fixed modification on cysteines. The mass tolerance for precursor and fragment ions was 4.5 ppm. False discovery rate (FDR) for the identification protein and peptides was 1%. Intensity-based absolute quantification (IBAQ) values were recorded. Reverse hits and contaminants were excluded.The sum of all IBAQ values of data sets were normalized to the corresponding control set. Protein abundance within native lanes were normalized to maximum appearance to enable comparison of mitochondrial complexes between control and mutant. All primary data were made publically available by the ProteomeXchange Consortium via the PRIDE [62] partner repository with the dataset identifiers PXD035352 (testis), PXD036901 (brain) and PXD036933 (heart). The entries contain the individual peptide sequences, single peptide identifications, all modifications observed, peptide identification scores, accession numbers, peptide assignments and quantification measurements.

### 2.7. Complexome profiling for OXPHOS subunits and other mitochondrial assemblies

Slice number of the maximum appearance of mitochondrial CI (979,577 Da), CII (122,945 Da) CIII dimer (483,272 Da), CIV (213,172 Da), CV (537,939 Da) and respiratory supercomplex containing CI, III dimer and one copy of CIV (1,676,021Da) was used for native mass calibration. The software NOVA (v.0.5.7) was used for hierarchical clustering of complexomics data [63].

### 2.8. Protein-protein-interaction bioinformatics and pathway enrichment analysis by STRING

To survey the known protein associations of the poorly studied AAA+ ATPase, data were extracted from the biomedical interaction repository BioGrid (https://thebiogrid.org/, last accessed on 8 November 2021), where endogenous VWA8 was detected in a complex with bait proteins. The STRING webserver (https://string-db.org/, last accessed on 8 November 2022) was used for visualization and statistics on these protein–protein interactions and pathway analyses. Filtered gene symbols were entered into the Multiple Proteins option for *Mus musculus*, and the graphical output was archived, highlighting factors with identical features by coloring, as explained in the figure caption.

### 2.9. Heavy metal analysis

Heavy metal abundance was measured in brain hemispheres (10 WT versus 10 CLPP-null) of 4-month-old mice as described before [64]. In brief, frozen mouse brain samples were thawed at room temperature, and transferred in total into a digestion tube. Dry matter was determined after oven-drying at 104 °C for 4 h. Due to the small amount of material, the dry matter could only be determined as a single measurement. Dried samples were subsequently mineralized through microwave digestion (Mars6, CEM GmbH, Kamp-Lintfort, Germany) using 3 ml of nitric acid, 1.5 ml of hydrogen peroxide and 2.5 ml water. Digested samples were analyzed for Ca, Co, Cu, Fe, Mg, Mn, Mo, Se, Vn, and Zn by inductively coupled plasma mass spectrometry (ICP-MS; PerkinElmer, Waltham, MA, USA), whereby indium and germanium were used as internal standards and commercial standard solutions for calibration.

### 2.10. Quantitative immunoblots

Protein was isolated with RIPA buffer (50 M Tris/HCl pH 8.0, 150 mM NaCl, 0.1% SDS, 1% Triton X-100, 0.5% sodium deoxycholate, 2 mM EDTA, and protease inhibitor cocktail from Sigma Aldrich). Protein content was determined using BCA assay (Life Technologies). 15 µg of protein were loaded for quantitative immunoblotting. Subcellular fractionations in mouse embryonic fibroblasts (MEF) (3 WT versus 3 *Clpp*-null)) were obtained as described before [65]. Subcellular fractionations from tissues were done as described in [66]. Primary antibodies used were as follows: CLPP (Proteintech, 15698-1-AP, 1:1000), CLPX (Abcam, ab168338, 1:1000), VWA8 (Invitrogen, PA5-58648, 1:1000), PLPBP (Proteintech, 25154-1-AP, 1:1000), HSPA9 (Oxford Biomedical Research, GR 02, 1:1000), TRAP1 (Abcam, ab109323, 1:1000), COX15 (Proteintech, 11441-1-AP, 1:1000), PTCD1 (Abclonal, A16219, 1:1000), MRPL38 (Proteintech, 15913-1-AP, 1:1000), MRPL18 (Novus, NBP2-94482, 1:1000), GFM1 (Proteintech, 14274-1-AP, 1:1000), OAT (Invitrogen, PA5-92842, 1:1000), ALDH18A1 (Proteintech, 17719-1-AP, 1:1000), HSP60 (SantaCruz sc-13115; 1:500), GAPDH (Calbiochem, CB1001; 1:1000). Secondary antibodies (1:15000) were fluorescence-labeled, obtained from Li-Cor, and either anti-mouse or anti-rabbit. The fluorescent signals were detected by the Li-Cor Odyssey Classic Instrument and was densitometrically analyzed with Image Studio Lite version 5.2 (Li-Cor Biosciences).

### 2.11. RT-qPCR

Total RNA was isolated from testis tissues (3 WT versus 3 *Clpp*-null)) with TRI reagent (Sigma-Aldrich), and reverse transcription was done with SuperScript IV VILO Master Mix (Thermo Fisher Scientific). Reverse-transcriptase real-time quantitative polymerase chain reaction (RT-qPCR) was carried out with TaqMan® Gene Expression Assays (Thermo Fisher Scientific) in complementary deoxyribonucleic acid (cDNA) from 10 ng total RNA in 10 μl reactions with 2× Master Mix (Roche) in a StepOnePlus Real-Time PCR System (Applied Biosystems). The data were analyzed with the 2^−ΔΔCT^ method [67]. The following Taqman assays were applied: *Aldh18a1*: Mm00444767_m1, *ClpP*: Mm00489940_m1, *ClpX:* Mm00488586_m1, *Cox15*: Mm00523096_m1, *Gfm1*: Mm00506856_m1, *Hspa9:* Mm00477716_g1, *Mrpl18*: Mm00775800_g1, *Mrpl38*: Mm00452473_m1, *Oat*: Mm00497544_m1, *Plpbp*: Mm00475293_m1, *Ptcd1*: Mm00505236_m1, *Sfxn4:* Mm00446462_m1, *Trap1*: Mm00446003_m1, *Tbp*: Mm00446973_m1, *Vwa8*: Mm01324241_m1.

### 2.12. Coimmunoprecipitation

Tissues were lysed in lysis buffer (20 mM Tris/HCl pH 8.0, 137 mM NaCl, 2 mM EDTA, 1% NP40, 1% glycerol with Protease-Inhibitor Cocktail cOmplete (Roche) by 30 min head-to-head rotation at 4 °C. Nuclear debris was removed by centrifugation at full speed at 4 °C for 20 min. Protein content was determined by BCA (Life Techologies). 1000-1500 µg of protein lysate were incubated with 5 µg of pull antibody and rotated for 2 h at room temperature (RT). In the meantime, 1.5 mg of Dynabeads (Thermo Fisher, #10004D) were washed 3× with PBS/T and added to the lysate/antibody solution. The mix was rotated head-to-head for 60 min at RT. After 5 washes with PBS, the tubes were fixed on a magnetic stand and the elution was done with 40 µl of 50 mM glycin, pH 2.8 and the eluate was mixed with loading buffer, boiled for 5 min at 90 °C and loaded for SDS electrophoresis. Antibodies used for CoIP were: CLPX (Invitrogen, PA5-79052), CLPX (Abcam, ab168338), CLPX (NSJ Bioreagents, RQ5618), CLPP (Proteintech, 15698-1-AP), MRPL38 (Invitrogen, PA5-118074), GFM1 (Proteintech, 14274-1-AP), OAT (Invitrogen, PA5-92842).

### 2.13. Statistics

Statistical analysis of quantitative immunoblot and RT-qPCR results were conducted by using GraphPad Prism (version 8.4.2, GraphPad) with unpaired Student’s t-tests with Welch’s correction. The results including standard error of the mean (SEM) and p-values were visualized in bar graphs, with the following significances illustrated by asterisks: * p < 0.05; ** p < 0.01; *** p < 0.001; **** p < 0.0001; T (statistical Trend) 0.05 < p < 0.1.

## 3. Results

To obtain complexomics profiles and judge their consistency, mitochondria from three tissues were purified via sucrose gradients before performing protein extraction with 20% digitonin detergent, blue-native electrophoresis (BNE), and label-free quantitative mass spectrometry of 48 gel slices. Subsequent analyses were performed to identify (1) abundance/accumulations of individual proteins under native conditions, their potential association/comigration within protein complexes, and their dispersion in mutant samples. The primary data from testis, brain, and heart are provided in Table S1, and were deposited publically at PRIDE. For validation experiments by quantitative immunoblots, mitochondria were isolated by differential detergent cell fractionation. Crucial findings were then assessed in separate samples via coimmunoprecipitation experiments.

### 3.1. Migration pattern and abundance of AAA+ proteins indicate CLPX and VWA8 both to depend strongly on CLPP

As expected, CLPP protein was absent from mutant samples and migrated in wildtype (WT) samples at positions that correspond to a homoheptameric ring (Figure 2). However, no CLPP peptides were detected where the CLPXP proteolytic barrel would run in BNE (expected molecular size around 565 kDa), and only 5% of CLPX comigrated with the homoheptameric CLPP ring in WT testis. Also unexpectedly, the abundance of endogenous CLPX, LONP1 and ATAD3 in WT tissues peaked at monomer to dimer positions, with no CLPX at all being detected at the expected homohexamer position. These surprising multimerization deficits may be explained by two alternative scenarios: (1) Our samples were frozen and treated with sucrose and digitonin, possibly resulting in the dissolution of relatively unstable complexes. However, blue native gels with samples that were never frozen also showed anti-CLPP immunoreactivity at the homoheptamer migration position of rather than the expected CLPXP proteolytic barrel. The proteolytic CLXP chamber appears only when CLPP cannot complete its function (see [45], Supplementary Figure EV3 B). (2) Alternatively, most CLPX may serve unfoldase-foldase-chaperone functions independent from CLPP degradation activity, for example assisting PLP to access the appropriate fold within ALAS for porphyrin biosynthesis. While we are studying endogenous mitochondrial proteins in the absence of degradation stress during slow growth, the previously published structural analysis of CLPXP barrels employed bacterial overexpressed and purified proteins, mixed at saturating conditions, from cells growing at logarithmic rates where protein degradation capacity may be maximized [68].

**Figure 2.**
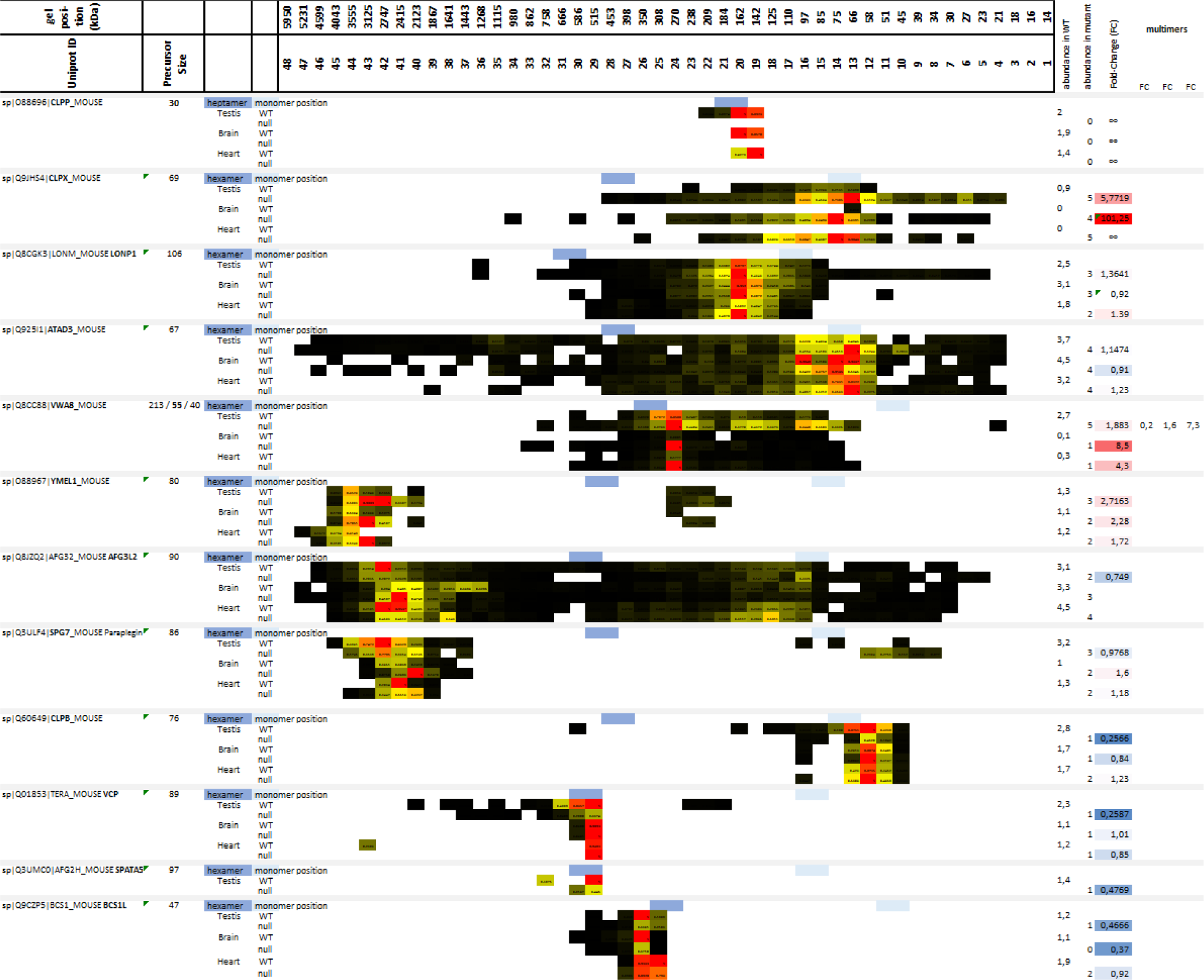
Analysis of mitochondrial AAA+ unfoldases in complexomics profile of WT versus CLPP-null mouse testis, brain and heart shows selective CLPX and VWA8 accumulation. Successive columns show the identity of each factor providing the UniProt-ID code, then molecular sizes of each precursor protein and potential isoforms according to UniProt database, tissue type, and the genotype for each line. Further columns illustrate the 48 gel slices representing protein complex sizes (from about 6,000 kDa on the left, until 10 kDa at the right), with white fields representing a lack of detection, while colored fields represent detection with max_norm values (dark brown for lowest, green for medium, bright red for highest abundance). Label-free mass spectrometry was used to quantify abundance in each slice, and the summed abundance across all slices was shown in digits on the right side, displaying mutant total, WT total, and the ratio as fold-change (FC) in a heatmap with strongest accumulations again in red, while strongest decreases are highlighted in blue. For VWA8, several isoforms are known to exist, and the FC was calculated for three separate migration peaks. Given that AAA+ unfoldases are thought to assembly in homohexamers, the expected migration position for each factor was derived and shown for the monomer (light blue fields) as well as the homohexamer (dark blue fields). Confirming the genotypes, CLPP was absent from all mutant tissues. Overall, CLPX, LONP1 and ATAD3 migrated much faster than expected for their homohexameric assemblies, while YMEL1, AFG3L2 and SPG7 migrated much slower than expected for homohexamers. CLPX and VWA8 showed consistent accumulation across the three tissues (LONP1 only in testis and heart) and migrated in more disperse pattern when CLPP was absent. In CLPP-null more than WT tissues, CLPP and VWA8 migration positions overlapped.

The observation that mature CLPX accumulated and migrated not only at its expected monomeric position around 60 kDa, but physiologically up to sizes around 250 kDa, and in CLPP-null tissues until sizes of 450 kDa, made it possible to use coaccumulation, comigration and dispersion patterns as criteria to identify potential CLPX interactors in the matrix. Overall, the scarcity of mitochondrial matrix proteins that comigrate with CLPX, as well as showing accumulated abundance upon CLPP deficiency, was very helpful to identify candidate interactions, which can be validated by further technical approaches.

Several migration positions existed for VWA8, and they were roughly compatible with dimer, trimer and homohexamer sizes for the medium mouse isoform around 55 kDa, but not for the small (40 kDa) and large (213 kDa) isoforms (Figure 2). The inner membrane-associated AAA+ proteases YME1L1, AFG3L2, SPG7 showed unexpectedly high migration sizes, comigrating around 3 MDa, consistent with reports that YME1L1 exists in a complex with PARL and SLP2 [69], while AFG3L2 and SPG7 are tethered to PHB1 [70] (Figure 2). Migration positions are compatible with the expected homohexameric assembly were observed only for CLPB, VCP, SPATA5 and BCS1L.

CLPP deficiency caused accumulation and dispersed migration in overlapping positions for the matrix unfoldases CLPX and VWA8 in all three tissues (Figure 2), the matrix unfoldase/protease LONP1 accumulation and dispersion in two tissues, and inner membrane unfoldase YME1L1 accumulation at much higher migration positions than CLPX (this metalloprotease acts on the respiratory complex IV [71] and in the in the mitochondrial intermembrane space, rather than the matrix), while the other mitochondrial AAA+ unfoldases appeared unchanged. Thus, only VWA8 shows similarity to CLPX in mutant tissue. Importantly, a meta-analysis of proteins that pulled VWA8 as prey in interaction assays according to the BIOGRID database (accessed on Nov 7, 2022) identified the mitochondrial RNA granule and mitoribosomal factors, as well as the respiratory chain complexes, as putative targets of VWA8 (Figure S1), in good overlap with previously reported CLPX targets [45, 49, 53, 56].

### 3.2. Comigration of mtSSU monomeric factors with ERAL1 as expected, but only mtLSU intermediate assemblies comigrate with CLPX and VWA8

To understand mechanistically how CLPXP targets the mitoribosomal translation machinery, the physiological and pathological migration range of target-selecting CLPX was now compared firstly to mitoribosomal proteins (Figure 3), then to other ribonucleoproteins (Figure S2), furthermore to PLP-associated proteins and additional chaperones (Figure S3), and finally to the mitoribosomally translated respiratory chain components (Figure S4).

**Figure 3.**
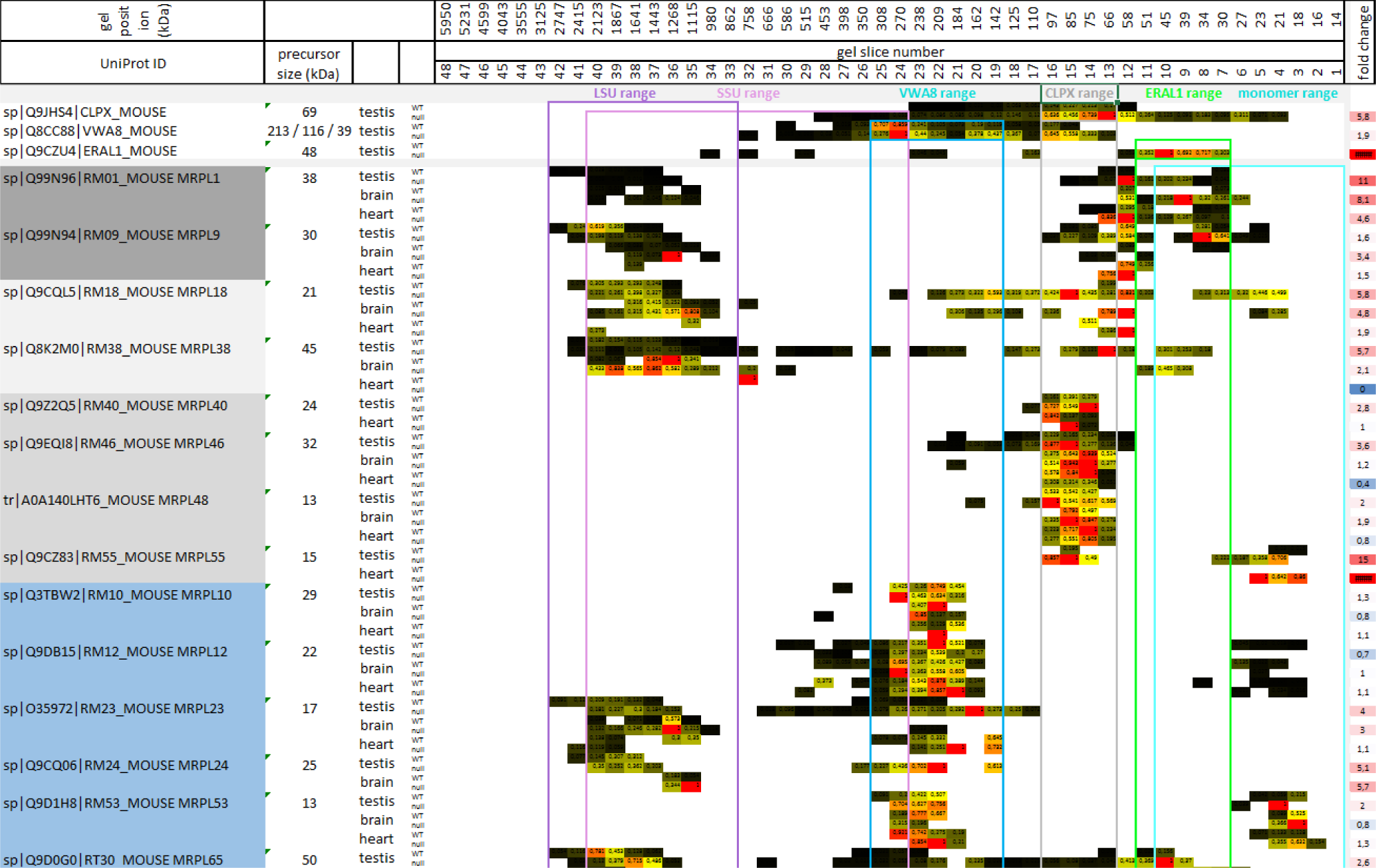
mtLSU factors in complexomics profile of WT versus CLPP-null mouse testis, brain and heart. To define possible interactions, migration patterns were compared with the detection range of AAA+ unfoldases CLPX (grey box) and VWA8 (dark blue box), as well as the mtSSU-folding rRNA chaperone ERAL1 (green box). For orientation, the range of monomeric mitoribosomal factors (light blue box), the assembled mtLSU (purple box) and the assembled mtSSU (lavender box) is shown. All detectable mtLSU and mtSSU complexomics data were compiled in Table S1. The format is analogous to Figure 2. Numbers at the right margin represent the null/WT total abundance ratio as fold-change (FC). The heatmap illustrates the strongest accumulations again in red, while the strongest decreases highlighted in blue. Overall, CLPX and VWA8 did not comigrate with fully assembled LSU or monomeric proteins, but instead with two different intermediate assemblies of the LSU. The ERAL1 detection range showed almost no overlap with the CLPX range in WT tissues.

Given a previous claim that the rRNA chaperone ERAL1 mediates the CLPP-null effect on mitoribosomes [45], it was surprising to find that the CLPX migration range in BNE gels had little overlap with the positions where ERAL1 was detectable (Figure 3). In agreement with previous reports that ERAL1 acts in the assembly of the mtSSU with its 12S rRNA [46, 47], ERAL1 migrated around 50 kDa in overlap with some mtSSU factors at monomeric positions (Figure 3 and Table S1).

In contrast, CLPX migration occurred at higher molecular weights than monomeric proteins and lower than completely assembled mtSSU/mtLSU complexes. This finding casts doubts on the proposed CLPXP role for functional mitoribosomes that were stalled during translation. Instead, CLPX comigrated with intermediate assemblies of specific mtLSU factors (see Figure 3, and also Graphical Abstract in Discussion chapter). Such mtLSU intermediate assemblies were observed in BNE gels before and represent functional clusters [72–77], so they do not seem artifacts of tissue freezing. Beyond their positions within the CLPX range, these mtLSU factors were also conspicuous for their relatively strong accumulation and their dispersed migration in CLPP-null tissue, and therefore are credible CLPX targets. They include the stalk factors MRPL1 (average accumulation 8-fold) and MRPL9 (2-fold), as well as a previously described [73–75] mitochondria-specific cluster among central protuberance components, namely MRPL40 (2-fold), MRPL46 (2-fold), MRPL48 (2-fold) and MRPL55 (also known as bL31m, accumulation 15-fold in testis, infinite in heart), presumably together with MRPL18 (4-fold) and MRPL38 (3-fold) (see Figure 3 in [78], Figure 4 and extended Figure 8 in [79], and [77]). The L1-stalk of the mtLSU has been implicated in tRNA translocation [80], while the L18-L38-L40-L46-L48 cluster serves to hold the mitochondrial tRNA-valine or tRNA-phenylalanine or 5S-rRNA in the central protuberance (see Figure 3 in [78], and [81]). It is interesting to note that MRPL18, MRPL25, MRPL46, MRPL48 and MRPL55 are not phylogenetically conserved in *Saccharomyces cerevisiae*, just like CLPP. Three proteins showed much higher migration positions in the CLPP-null tissues, namely MRPL18, MRPL38 and MRPL46, suggesting that tRNA-valine integration problems persist during ever larger assembly stages, and show a very similar dispersion/accumulation pattern in complexomics as CLPX. Given that CLPP loss triggers the identical phenotype in humans as mutations in the mitochondrial tRNA ligases HARS2 and LARS2, these observations may highlight an impact of CLPXP on the folding and assembly of tRNA with mtLSU central protuberance (CP). In contrast to the quite selective CLPP-null effect on only few mtLSU factors, non-selective accumulation was observed for practically all mtSSU factors, without CLPX comigration (Table S1), possibly due to mtSSU interaction with accumulated LRPPRC/SLIRP/HARS2/TARS2 plus tRNAs.

**Figure 4.**
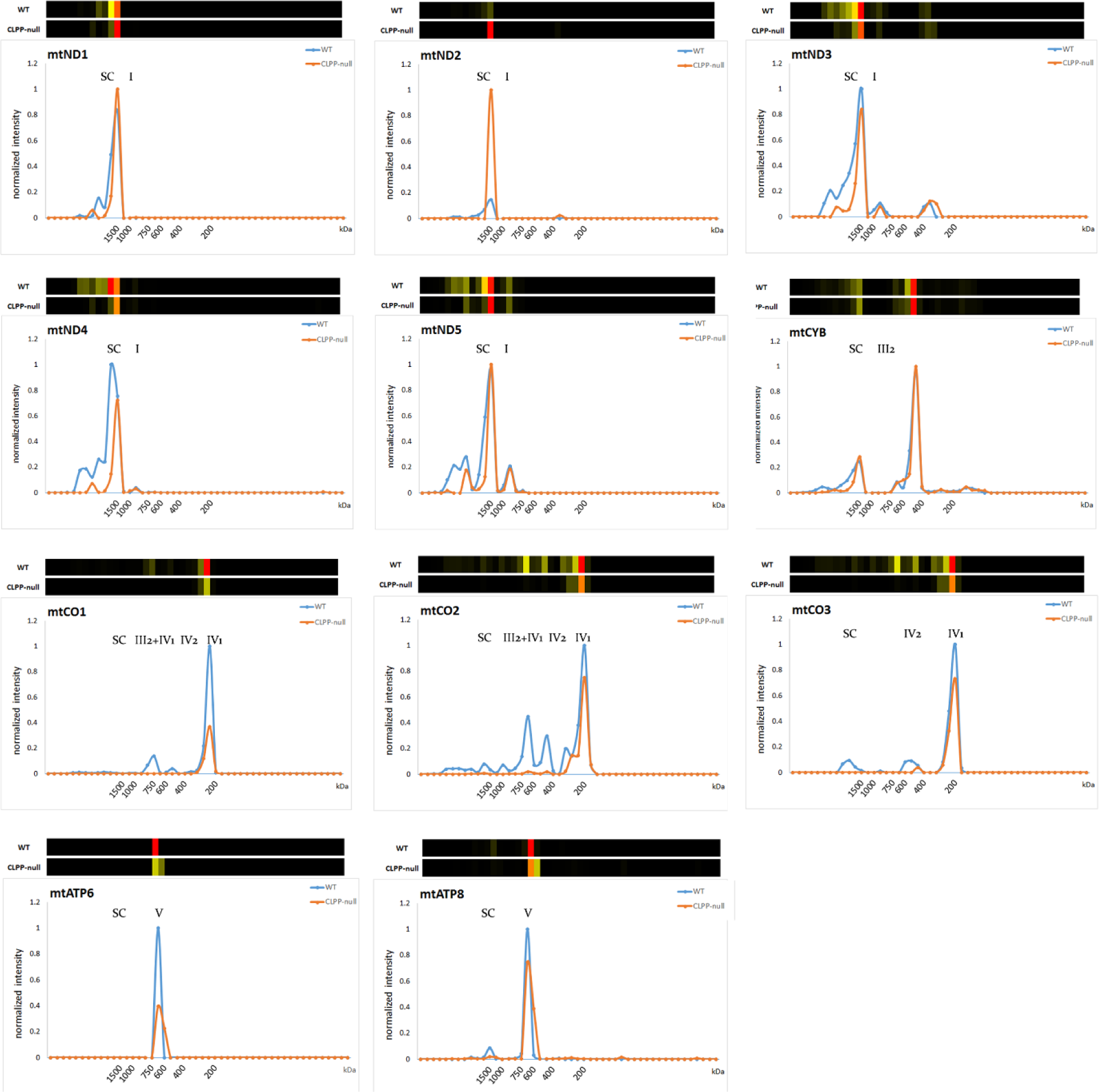
Migration profiles of mitochondrially-translated factors in testis as most severely affected tissue. Each subpanel shows the amount of a protein at different migration positions, above illustrating it as heat map, and below providing a plot of relative abundance on the Y-axis, with the X-axis representing the 48 slices of BNE gels with the scale of approximate molecular mass in kDa. The expected size of fully assembled monomeric complexes is denoted in each panel as Roman numeral, with Arabic numerals behind indicating homo-multimerization status (1 for monomer, 2 for dimer). Monomeric CI runs around 1 MDa, dimeric CIII around 0.5 MDa, monomeric CIV around 0.2 MDa, CV around 0.7 MDa, supercomplexes 0-4 (SC) in the range around 1.5 MDa and above [75, 108, 109]. Although not distinguished in the panels for space reasons, CI+CIV are known to associate, before the supercomplex S0 containing I + III_2_, and the supercomplex S1 containing I + III_2_ + IV are assembled [109]. The multimerization of CIV_1_ to CIV_2_ at about 0.45 MDa and its association as CIII_2_-CIV_1_ at about 0.7 MDa are known higher assemblies of MTCO1-2-3 [110]. The defect of homo-multimerization and association with other respiratory complexes was particularly severe for complex-IV, as seen for proteins MTCO1-MTCO2-MTCO3.

The VWA8 migration peak was around 200 kDa at higher molecular weights than the 80 kDa CLPX peak, while the two migration ranges overlapped. Again, VWA8 comigrated with a mtLSU intermediate assembly complex, suggesting that it acts during the association or dissociation stage rather than the repair of complete mitoribosomes. This is consistent with previous observations that the yeast VWA8 ortholog Midasin/Rea1 acts in the maturation/assembly of pre-mtLSU central protuberance and L1 stalk with 5S rRNA [82]. In mouse, VWA8 comigration concerned on the one hand the MPRL10-MRPL12-MRPL53 proteins as components of the LSU L7/L12 stalk (a subcomplex that was already reported and comigrates also with LRPPRC and SLIRP [72, 73]), but the CLPP-null mutation did not cause them to accumulate or migrate differently. The L7/L12 stalk is the docking site of translation elongation GTPases (orthologs of GFM1/2) [83]. On the other hand, the VWA8 comigration with the LSU NPET (nascent polypeptide exit tunnel) wall proteins MRPL23-MRPL24 [78] was accompanied by altered migration and accumulation (on average 3-fold for MRPL23, 5-fold for MRPL24). MRPL65 was previously named MRPS30, and its position in the latest mtLSU assembly is shown in [78] Figure 1 right panel. MRPL65 also showed VWA8 comigration that appeared altered upon CLPP-deficiency, and accumulation in the mutant (3-fold). Thus, when mitoribosomal translation fidelity is impaired in CLPP-null tissues due to CLPX-dependent folding problems at the mtLSU central protuberance, the unfoldase function of VWA8 might target specific NPET components in mtLSU disassembly intermediates.

Serendipitously, we observed the distribution of MRPS36/KGD4 to be well above the size of the completely assembled mitoribosome, arguing against its integral role for the mtSSU and supporting recent evidence that it is a subunit of the mitochondrial alpha-ketoglutarate dehydrogenase complex [84, 85].

All other ribonucleoproteins were assessed and 16 candidates were conspicuous in all three tissues (Figure S2), with translation factors occupying a prominent place. Among the disease proteins responsible for Perrault syndrome, HARS2 was prominent because of its strong accumulation in CLPP-null tissue, as previously reported upon quantitative immunoblots [17], and here the complexomics profile revealed a parallel distribution of CLPX and HARS2 from their monomeric sizes (precursors of 69 and 23 kDa, respectively) until molecular sizes around 250 kDa. Similarly, a close comigration with CLPX was documented for GFM1 from its monomeric size (84 kDa) to positions around 250 kDa, in addition to very strong accumulation (average 10-fold). A recent review of putative CLPXP substrates in different species concluded that the translation elongation GTPase GFM1/GFM2 orthologs are exceptionally consistent candidates [49]. Curiously, the extra-mitochondrial translation initiation complex core factor EIF3C (precursor size 106 kDa) also appeared in CLPP-null mitochondria greatly accumulated (average 5-fold) and showed comigration across the entire CLPX distribution range. EIF3C is a core factor in the EIF3 complex that recruits tRNA-methionine during pre-initiation to scan for start codon versus internal ribosomal entry sites under stress conditions [86–89].

In addition, several proteins with localization at the mitochondrial RNA granule were co-migrating with CLPX. Moderate accumulation in CLPP-null tissue (LRPPRC 3-fold, SLIRP 12-fold) but clear dispersion was observed for both components of the LRPPRC-SLIRP complex, which associates preferentially with the 5’-untranslated region of *mtCo1* and *mtCo3* mRNA, delivering them to the mRNA channel in the mtSSU via PTCD3/MRPS39 [90], and is crucial for isolated complex-IV deficiency [91–94]. While they showed only minor overlap with the physiological CLPX migration range, their migration position coincided with the high-molecular weight positions reached by accumulated CLPX in the mutant, and coincided with the VWA8 migration range. Much stronger accumulation (ratios until infinite values) and CLPX/VWA8-comigration/dispersion were documented for additional RNA-binding PPR-domain-containing factors such as PTCD1 (and PTCD2) as pseudouridylation enzyme, which processes rRNA before integration into the mtLSU [95], but also performs processing of tRNA-ligase for leucine (LARS2) and controls respiratory complex IV activity [96, 97]. Given that mutations in the mitochondrial leucine tRNA-ligase LARS2 cause the Perrault syndrome just like CLPP deficiency [98], this observation may further strengthen the notion of RNA processing importance in this pathogenesis. Similar massive upregulation and CLPX-comigration/dispersion were also apparent for the G-quadruplex RNA unwinding factor GRSF1, which mediates protective effects against iron overload and ferroptosis [99], and which interacts with the mitochondrial RNase P complex subunit TRMT10C/MRPP1 [100]. Modest accumulations were also found for PLP-dependent SHMT2 and MTHFD2, which are needed for tRNA methylation in mitochondria [101, 102], otherwise stalled mitoribosomes would occur [103]. Jointly, these observations suggest that specific mitoribosomal translation factors may be the main physiological targets of CLPXP, but particularly upon CLPP deficiency, the excess CLPX/VWA8 may also target RNA granule components.

Accumulation, CLPX/VWA8-comigration, with dispersion in CLPP-null tissues was also documented for various molecular chaperones, and enzymes whose activity is modulated by the chaperone PLP (Figure S3). Importantly, the Pyridoxal Phosphate Binding Protein (PLPBP or PROSC) showed migration positions until 70 kDa in mutant brain instead of its WT position around 30-35 kDa. As expected consequence of the gain-of-function of CLPX and the PLP, their established target ALAS2 as rate-limiting enzyme of porphyrin/heme biosynthesis in most tissues showed massive accumulation. Strong accumulation, dispersion and comigration with CLPX was apparent also for the PLP-dependent enzyme OAT, and similar accumulation concerned also the enzyme ALDH18A1, as previously reported in global proteome profiles [56, 104]. Both factors play connected pathway functions in the interconversion of amino acids such as proline, glutamate and arginine. Smaller accumulations and some dispersion were observed for the pyruvate homeostasis enzymes PDK1, PDK3 and PDPR.

In view of the phylogenetically conserved interaction between CLPX and HSP70 family members in the chaperone pathway, it was no surprise to find accumulations of HSPA9 (Mortalin) and the mitochondrial HSP70-cochaperones/modulators GRPEL1-GRPEL2-DNLZ-CCDC127 [105–107], but also TRAP1 as mitochondrial member of the HSP90 family showed similar accumulation. However, both chaperones have molecular sizes that are very similar to the CLPX precursor around 69 kDa, so their comigration would be coincidence. Overall, the loss of CLPP peptidase and accumulation of CLPX unfoldase has a profound impact on factors that interact with the chaperone PLP and determine folding/assembly.

### 3.3. CLPP-null OXPHOS complexes affected more by assembly than translation problems

The CLPXP complex has a prominent impact on mitoribosomal translation, which is responsible for the biosynthesis of 13 well-characterized proteins that have key roles in the respiratory chain complexes CI, CIII, CIV and CV. While these factors are embedded in the mitochondrial inner membrane, many regulatory factors, assembly factors, and factors that assist in the flow of electrons are assembled around them in complexes that protrude in the matrix space. This matrix immersion is particularly true for the N-module of CI, and indeed the three core proteins of the N-module (NDUFS1, NDUFV1, NUDFV2) were shown to be CLPP-dependent in their exceptionally high turnover rate [53]. Analyzing the complexome profiles regarding these N-module proteins (Figure S4 top), they were found to comigrate with CLPX across a wide range in WT samples, and even more in CLPP-null tissues. Interestingly, the abundance of NDUFS1-NDUFV1-NDUFV2 was increased selectively in CLPP-null slices where they comigrated with CLPX (Figure S4 top, fold-changes at right margin), suggesting inefficiency and slowed turnover during assembly interactions. In contrast, their abundance was decreased in the slices that represent fully assembled complex-I, suggesting that their integration into their N-module position is impaired. Altogether, these complexome data confirm a role of CLPXP for the CI N-module core factor, but apparently regarding their assembly/disassembly rates rather than their degradation.

The complexomics profile of mitochondrially encoded/translated factors in the respiratory chain again showed lowered abundance rather than accumulation in CLPP-null tissue, in good agreement with the notion of impaired mitoribosomal translation fidelity. Prominently strong reductions were found in testis, as the most severely affected tissue of CLPP-null mice that model the complete infertility of PRLTS3 patients. The reductions reached levels of 0.36-fold for MTND4, 0.32-fold for MTCO1 (COX1), and 0.36-fold for MTCO2 (COX2) (Figure S4 middle, right margin).

However, deficits of assembly seemed much more severe than deficits of translation. As shown in Figure 4, monomeric complexes I-III-IV-V formed quite adequately in CLPP-null testis as the most severely affected tissue, but the multimerization steps to form supercomplexes (SC) were generally deficient. Almost absent integration into higher-order SC was observed for MTND3-MTND4-MTND5, and for MTCO1-MTCO2-MTCO3 practically no assembly into the CIV_2_ dimer, or the CIII_2_-CIV_3_ intermediate assembly, or SC 0-4 was detected (Figure 4) (mutant testis showing fold-changes of 0, 0.04, and 0, respectively, in the migration range from IV_2_ until SC). It is therefore interesting to note that in WT tissue, VWA8 comigrated with the CI Q-module assembly factor NDUFAF3 and ND1-module assembly factor TIMMDC1, as well as the mitochondrial CI intermediate assembly (MCIA) complex members TMEM186, NDUFAF1, ECSIT, ACAD9, TMEM126B that are responsible of the ND2 module, all of which accumulated in CLPP-null testis (Table S1). An uncommonly strong 17-fold accumulation together with a migration shift to a much higher, possibly dimeric/trimeric position was noted in CLPP-null brain for SFXN4, which was recently identified to modulate iron homeostasis and act as MCIA component for CI assembly [111]. Accumulation (2.7-fold) was also documented for SLC25A28 as mitochondrial iron transporter upon CLPP depletion, but this was detectable only in testis tissue.

Furthermore, comigration with CLPX in WT tissue was observed for practically all components of respiratory CIV (Figure S4). As a CIV component protein, which showed outstanding accumulation in CLPP-null tissues (in testis 8-fold) and comigrated with CLPX in its most abundant positions (slices 12-16), the complexome profiles identified COX15 (Figure S4 bottom). This protein is not a structural constituent of the fully assembled complex-IV, but is essential for its biogenesis and assembly. Curiously, COX15 is not immersed in the matrix, but has five transmembrane domains that localize it as homo-oligomer in the mitochondrial inner membrane [112, 113]. Thus, it is unlikely to be a direct target of CLPXP functions, but its selective strong affection might be indirect: CLPP-null mice are known to have elevated hemoglobin [114] and probably also heme levels, and COX15 uses two heme-O to generate heme-A molecules that are needed as cofactors to mediate CIV assembly and activity [112, 115].

Overall, the impact of CLPXP on the respiratory chain appears strongest on CIV and CI, causing a deficit in total levels rather than the accumulation seen for mitoribosomal translation factors, but respiratory chain SC assembly problems were stronger than peptide chain synthesis problems, and may be secondary to the iron/heme homeostasis alteration via SFXN4-MCIA for CI, and via COX15 for CIV.

### 3.4. Quantification of heavy metals in CLPP-null tissue demonstrates significant increases in iron, molybdenum, cobalt and manganese

In view of the known role of excess CLPX for ALAS overactivity for heme biosynthesis, and of the accumulation of iron-associated SFXN4-COX15 as potential explanation for respiratory complex assembly deficits, the question arose if iron and other heavy metals as well as heme are dysregulated and enhance PRLTS3 pathology in nervous circuits. Deposition of excess iron was documented in many neurodegenerative disorders [116]. Postmitotic neurons in brain tissue may be particularly vulnerable to the chronic impact of heavy metal toxicity via reactive oxygen species, ferroptosis and inflammation. The iron analysis was extended to a survey of other heavy metals (see Table S2), since excess protoporphyrin is toxic and may chelate also magnesium, zinc, copper, nickel, vanadium, cobalt and manganese [117–122].

As summarized in Table 1, 40-20% increases were observed for iron, molybdenum, cobalt, and manganese. These data are consistent with the overactivity of heme production by ALAS, and indicate a more widespread metal pathology. The tissue excess of molybdenum and cobalt suggests increased levels of the pterin-based molybdenum cofactor (Moco) [123] and of corrin-ring-based cobalamine (vitamin B12), respectively, while the excess of manganese is compatible with maximal activity of Mn-containing enzymes such as arginase or mitochondrial superoxide-dismutase. Neurotoxicity is well established for manganese overdosage [124], while molybdenum toxicity represses growth and fertility [125]. Overall, the elevation of iron and several other heavy metals in CLPP-null tissue may contribute to the impaired homeostasis of heme and downstream heme-dependent assemblies of respiratory CI and CIV.

**Table 1.**
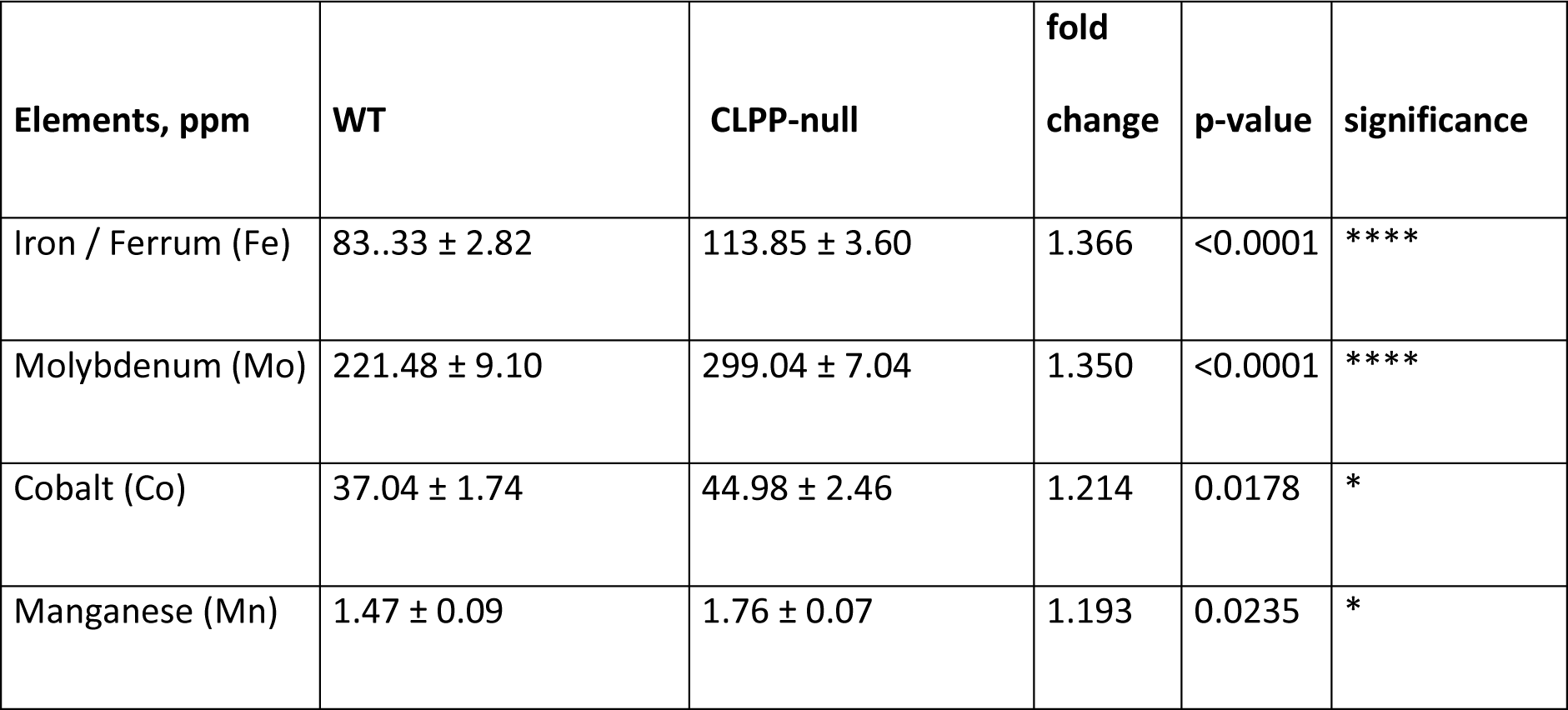
Accumulation of several heavy metals in CLPP-null brain. Quantification was done in 10 WT versus 10 CLPP-null brains at the age of 4 months by ICP-MS. Values are given in mean ± standard error per dry matter of tissue.

| Elements, ppm | WT | CLPP-null | fold change | p-value | significance |
| --- | --- | --- | --- | --- | --- |
| Iron / Ferrum (Fe) | 83.33 ± 2.82 | 113.85 ± 3.60 | 1.366 | <0.0001 | **** |
| Molybdenum (Mo) | 221.48 ± 9.10 | 299.04 ± 7.04 | 1.350 | <0.0001 | **** |
| Cobalt (Co) | 37.04 ± 1.74 | 44.98 ± 2.46 | 1.214 | 0.0178 | * |
| Manganese (Mn) | 1.47 ± 0.09 | 1.76 ± 0.07 | 1.193 | 0.0235 | * |

### 3.5. Validated accumulation of VWA8, CLPX, PLPBP, GFM1, MRPL18, MRPL38, HSPA9, TRAP1, COX15, PTCD1, ALDH18A1 and OAT in quantitative immunoblots

While the complexome profiles above assessed consistency between different tissues for all abundant mitochondrial proteins, subsequent validation experiments by quantitative immunoblots focused on relevant factors and tested their anomalies statistically, using several samples from testis or murine embryonic fibroblasts (MEF) depending on variant abundances and cross-reacting bands of both tissue expression patterns. For these experiments, mitochondrial fractions were isolated by differential detergents from cytosolic and nuclear fractions, and specificity and sensitivity analysis was performed for antibodies that were later used in coimmunoprecipitation studies. Convincing detection of endogenous protein levels (see Figure 5A) was achieved for CLPX and VWA8 as AAA+ unfoldases, PLPBP as PLP-chaperone homeostasis protein, HSPA9 and TRAP1 as molecular chaperones, COX15 as heme-A synthase for CIV assembly, PTCD1 as RNA granule component, MRPL38 and MRPL18 as mtLSU members, GFM1 as translation elongation factor and OAT as PLP-dependent amino acid homeostasis enzyme.

**Figure 5.**
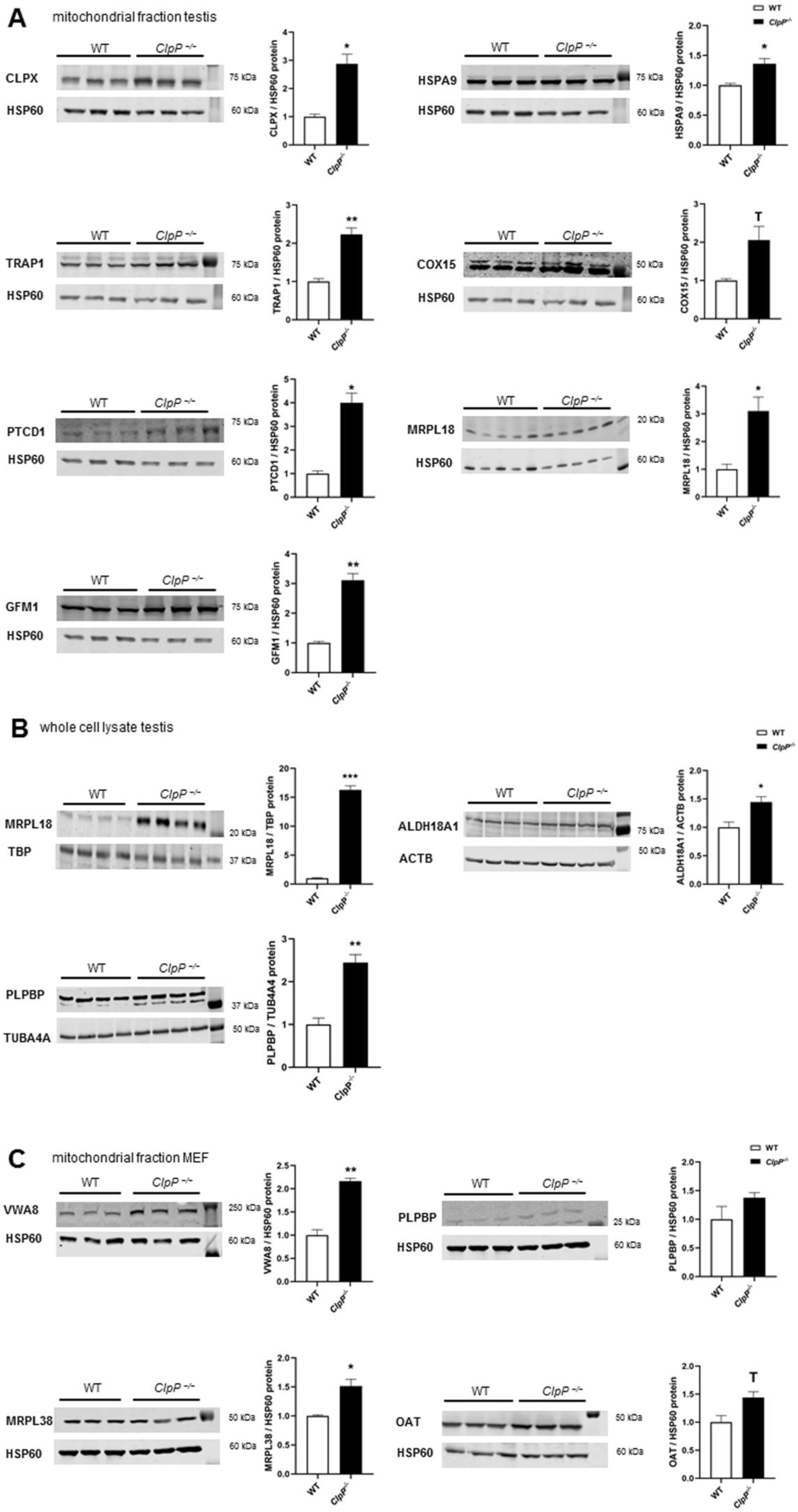
Experiments to assess mass-spectrometry abundance data by quantitative immunoblots. **(A)** Quantitative immunoblots in P21 testis subjected to differential detergent fractionation using the mitochondrial protein HSP60 as loading control to normalize abundance. **(B)** Quantitative immunoblots in 5-month-old testis tissue, whole cell lysates. Protein abundance levels were normalized against TBP, ACTB or TUB4A4 levels. **(C)** Quantitative immunoblots in MEF cells, subjected to differential detergent fractionation with mitochondrial HSP60 as normalizing protein. * p < 0.05; ** p < 0.01; *** p < 0.001; T (statistical Trend) 0.05 < p < 0.1.

In mitochondrial fractions from testis at postnatal day 21 (P21) where spermatogenesis has started but apoptotic cell loss is not yet noticed in mutants, using HSP60 as fraction marker and loading normalizer, accumulations were confirmed for CLPX (2.9-fold. p=0.003), HSPA9 (1.4-fold, p=0.04), TRAP1 (2.2-fold, p=0.01), COX15 (2.1-fold, p=0.10), PTCD1 (4.0-fold, p=0.01), MRPL18 (3.1-fold, p=0.02), GFM1 (3.1-fold, p=0.01). Solubilizing more protein via the SDS detergent in RIPA buffer, lysates of total testis tissue showed an even stronger accumulation of MRPL18 (16.3-fold; p=0.0002) and detected also significant accumulations of ALDH18A1 (1.4-fold; p=0.02) and of PLPBP (2.45-fold; p=0.001). In mitochondrial fractions of MEF, accumulations were replicated for VWA8 (2.16-fold, p=0.003), MRPL38 (1.5-fold, p=0.04), OAT (1.4-fold, p=0.05), but without significance for PLPBP (1.4-fold, p=0.23) in these cells in contrast to the severely affected CLPP-null testis. Overall, the quantitative immunoblots validated the principal complexome findings.

### 3.6. RT-qPCR analysis of transcriptional regulations

To understand if protein accumulations are due to delayed turnover as consequence of CLPXP proteolysis deficits, or due to transcriptional upregulation as possible compensatory effort, the expression of individual key factors was quantified by Reverse Transcriptase quantitative Polymerase Chain Reaction (RT-qPCR). Expression upregulations of *Hars2* and *mtCo1* in testis were reported previously [51]. To now elucidate the sequence of pathogenesis events, testis tissue was studied at postnatal days P17 and P21 (pre-manifestation) as well as P27 (post-manifestation). At the P27 stage, many transcripts show dysregulated levels, and it has to be taken into account that selective cell losses may account for some of the changes due to the rapid loss of spermatids at the end of meiosis-I. As shown in Figure 6, *Clpp* transcript was absent in all CLPP-null samples as expected, and *Clpx* transcript was downregulated at P27, presumably as cellular effort to compensate the pathological CLPX protein accumulation. Conversely however, *Vwa8* transcript was upregulated, suggesting that VWA8 protein accumulation serves to achieve some folding/assembly task that was inadequately performed by mutant CLPXP. Importantly, significant upregulations of *Aldh18a1* mRNA was observed already at P17, P21 and P27 (2.0-fold at p=0.03, 4.7-fold at p=0.02, 6.1-fold at p=0.002, respectively). Also *Oat* transcript seemed consistently elevated at P17 (1.65x, p=0.21), P21 (1.81x, p=0.22) and P27 (3.48x, p=0.002), but this became significant later than for *Aldh18a1*. These findings suggest that cells since pre-manifest stages maximize metabolic capacity of these enzymes to counteract their insufficient activity or/and impaired folding/assembly. At later post-manifest P27 in testis, similar transcript upregulations were also observed for translation factors (*Gfm1*, *Mrpl38*) and respiratory complex assembly factors (*Cox15*). Interestingly, transcripts of the iron-binding CI assembly factor *Sfxn4* seemed already increased at P17 (1.84x, p=0.20), with significantly upregulated mRNA levels at P21 (1.26x, p=0.03) and P27 (2.17x, p=0.04). The transcript of PLP-binding protein *Plpbp* was also elevated at P27 (1.8x, p=0.003), reflecting the complexome protein abundance results.

**Figure 6:**
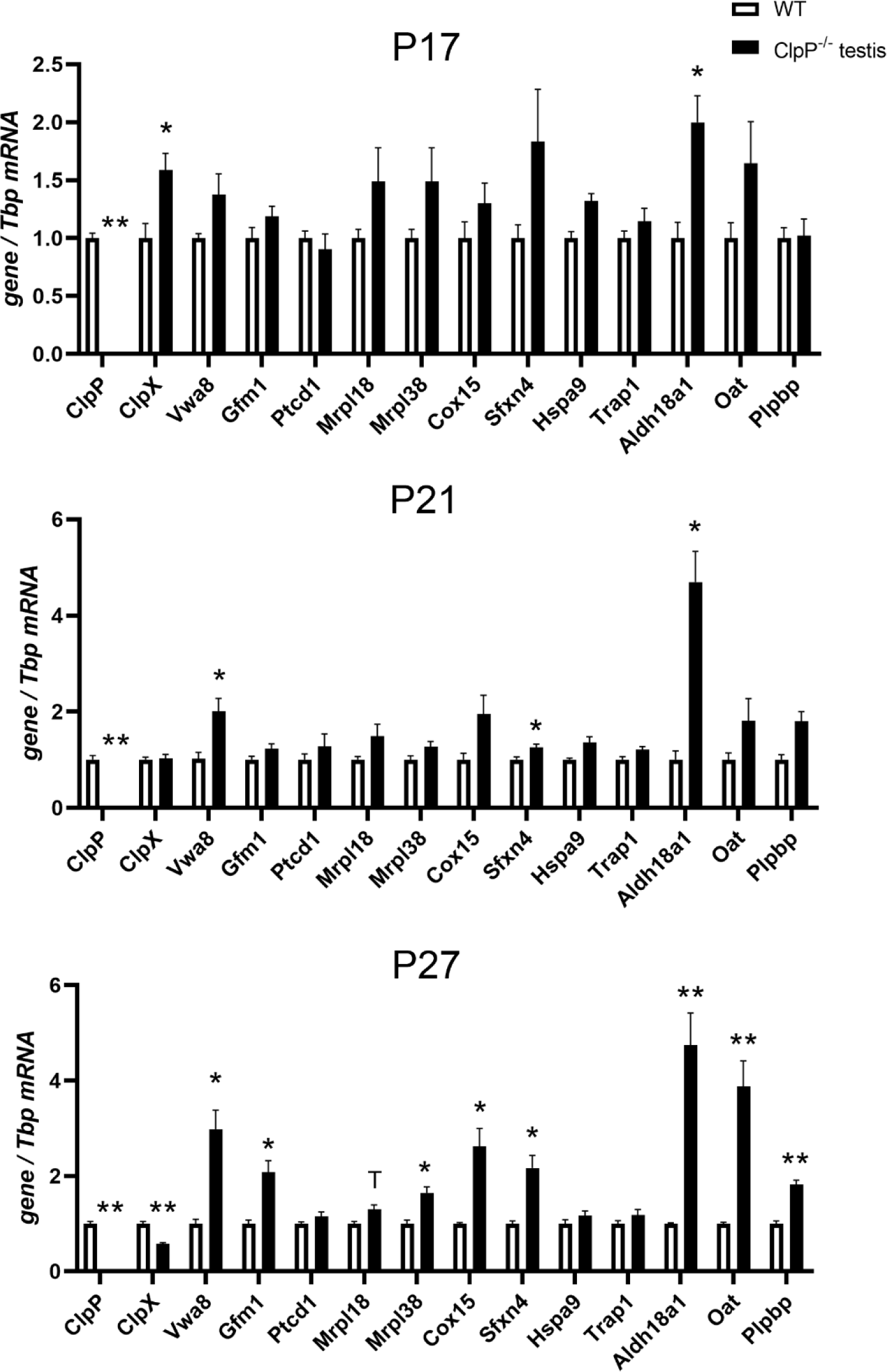
RT-qPCR studies in P21 and P27 testis. used *Tbp* mRNA levels as loading controls to normalize expression. * p < 0.05; ** p < 0.01; T (statistical trend) 0.05 < p < 0.1.

### 3.7. Coimmunoprecipitation of CLPX confirms interaction with MRPL18, GFM1 and OAT

To validate associations of CLPX with target proteins that were suggested by complexomics data, coimmunoprecipitation (CoIP) of CLPX and detection of endogenous protein complexes in tissue would be optimal. However, this approach depends firstly on the sufficient abundance of target proteins, secondly on the high-quality of antibodies for precipitation and detection, and thirdly on finding appropriate detergent/salt conditions. The CoIP conditions have to disrupt membranes of cells/mitochondria and dissociate mitoribosomes or respiratory chains sufficiently to permit the purification of interaction partners on beads, but these conditions must not interfere with the stable binding between CLPX and its substrate. We concentrated the validation work on GFM1 as translation elongation GTPase, the MRPL18/MRPL38 cluster as mtLSU-CP components that are folded around tRNA-valine or 5S-rRNA, and the amino acid homeostasis enzyme OAT which is soluble in the mitochondrial matrix. All of them were reported as targets of PLP binding [126–129]. The abundance of all these endogenous proteins is usually too low for detection in CoIP experiments, barely reaching the detection threshold in testis and liver where levels are maximal according to The Human Protein Atlas webserver (https://www.proteinatlas.org/ENSG00000166855-CLPX/tissue, last accessed on Dec 12, 2022). As expected however, the specific CLPX-MRPL38-GFM1-OAT immunoblot signals became reproducibly visible in CLPP-null tissue.

As shown in Figure 7, we prioritized GFM1 which was the most consistent CLPXP target among all surveys in various organisms [49]. Anti-CLPX antibodies (Invitrogen) coimmunoprecipitated GFM1 (detection by Proteintech anti-GFM1) from liver mitochondrial fractions. Conversely, upon CoIP of GFM1 (Proteintech) a CLPX band (detection by Abcam anti-CLPX) of expected size was detectable in CLPP-null liver whose cell lysates.

**Figure 7.**
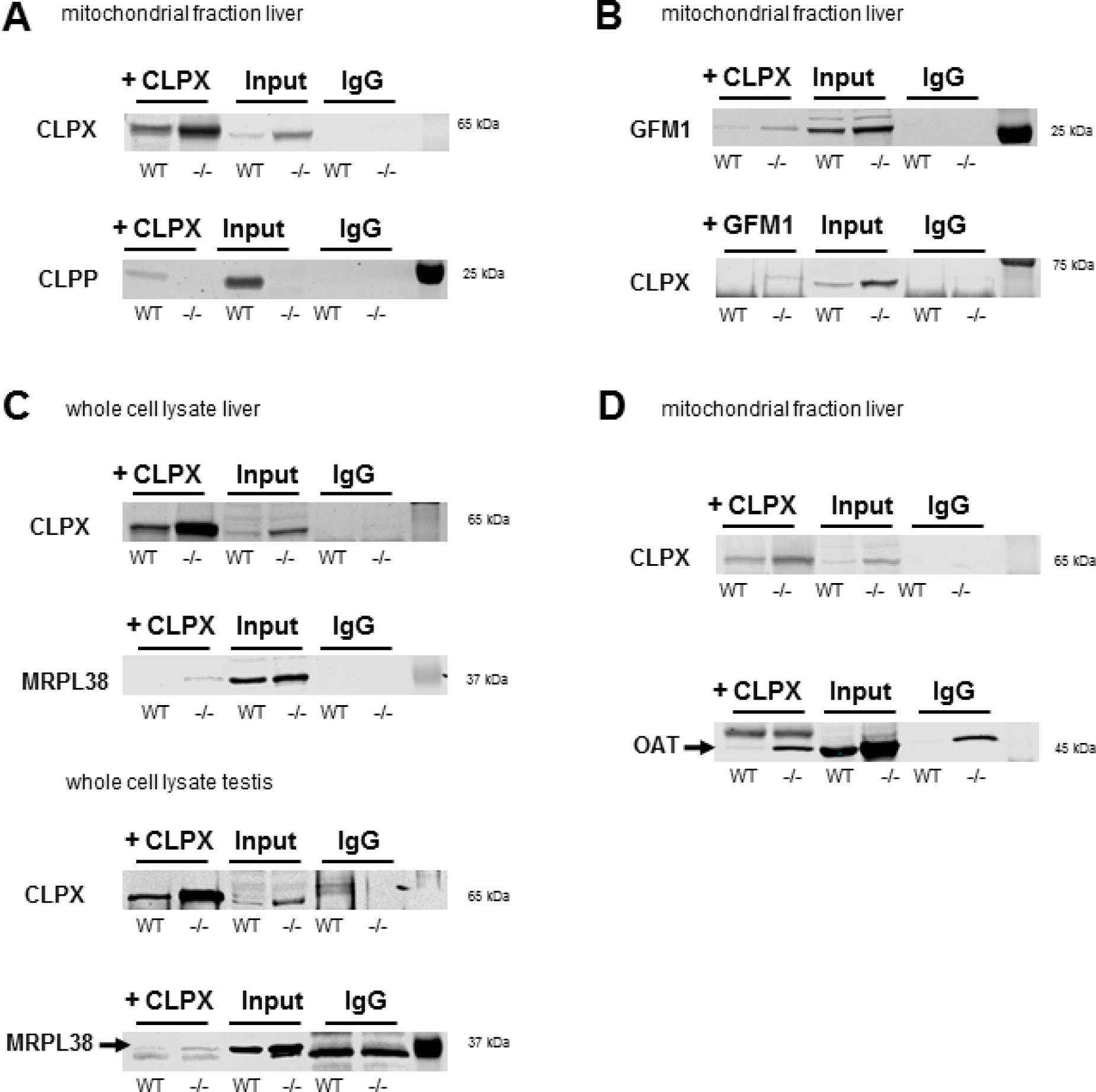
Experiments to validate mass-spectrometry complexomic by coimmunoprecipitation. The precipitation target protein is highlighted by the + sign. **(A)** Validation of IP by pulling with CLPX-antibody from Invitrogen, detecting CLPX with antibody from Abcam. Validation of CLPX-coimmunoprecipitation using CLPP (anti-CLPP antibody from Proteintech) as positive control of interaction, and assessing correct genotyping of CLPP-null tissues. **(B)** CoIP analyses in liver mitochondrial fractions used anti-CLPX (Invitrogen) and anti-GFM1 (Proteintech) antibodies with sufficient sensitivity and specificity to detect endogenous proteins, pulling down each of the two interaction partners as bait to assess the association with its prey. When the endogenous abundance of these proteins was at the threshold of detection in WT cells, their pathological accumulation in CLPP-null tissues made it possible to demonstrate associations. **(C)** CoIP analyses in whole cell liver and testis lysates, pulling with anti-CLPX (NSJ Bioreagents) and detection of CLPX (Abcam) as control gave also specific bands for MRPL38 (Invitrogen) in both tissues. In testis tissue, a slightly lower unspecific band appeared in the IgG control samples, the height of the correct band is marked with an arrow. **(D)** CoIP analyses in liver mitochondrial fractions pulled with anti-CLPX (Invitrogen) and confirming the IP with anti-CLPX (Abcam) showed specific bands for OAT (Invitrogen). A slightly higher unspecific band was detected in CLPP-null IgG control sample, the correct height of the band is marked with an arrow.

Given that the antibody against MRPL18 was of insufficient specificity for CoIP, MRPL38 as its closest interaction partner with tRNA-valine was analyzed. Again, the CoIP of CLPX (NSJ anti-mouse-CLPX) was able to pull MRPL38 in CLPP-null liver whole cell lysate more than WT (detection by Invitrogen anti-MRPL38), as well as in WT and mutant testis whole cell lysate (detection by Invitrogen anti-MRPL38), as demonstrated by appearance of a specific band with correct size and stronger abundance in the CLPP-null sample. Furthermore, the CoIP of CLPX (Invitrogen) was also able to pull OAT (detection by Invitrogen anti-OAT) again from CLPP-null liver mitochondrial fraction. While the antibodies against MRPL38 and OAT were specific enough for immunoblot detections, they did not have the quality to be used in the converse CoIP of MRPL38 and OAT to detect associated CLPX.

Although the CoIP data in WT tissue could not always unequivocally demonstrate physiological interactions between the two endogenous proteins, our CoIP findings in the pathological CLPP-null tissues coincide with previously published overexpression experiments and may underlie PRLTS pathology. Overall, these findings provide evidence that CLPX modulates the access of PLP not only to ALAS and thus modifies folding/activity/stability of this target protein, but similarly interacts also with GFM1 / mtLSU-CP proteins / OAT as known targets of PLP.

## 4. Discussion

The targets of CLPXP proteolytic activity remain controversial, despite analyses over decades in numerous organisms with diverse technical approaches, most of which involved recombinant overexpression [49, 55]. As a novel approach, we now focused on endogenous proteins in their stable interactions within three mouse tissues, trying to define potential CLPX targets based on four criteria: (1) their comigration with CLPX in complexome profiles of WT tissues, (2) their coaccumulation with CLPX / VWA8 after CLPP depletion, (3) the appearance at higher molecular weight positions in overlap with CLPX or VWA8 positions after CLPP depletion, (4) the consistency of such findings between testis, brain, and heart. Preliminary validation experiments confirmed CLPXP dependence for a few selected findings such as GFM1 accumulation, as proof-of-principle that this approach provides valid insights. Within the unbiased complexome profiles of mitochondria, we paid particular attention to the impact of CLPXP (i) on other AAA+ unfoldases to explore their functional overlap, (ii) on the RNA processing and translation machinery given previous evidence from bacteria to mice [45, 49, 53, 56, 130–132], (iii) on mitochondrially translated components of the respiratory chain, and (iv) on matrix enzymes, attempting to elucidate the molecular anomalies at these sites of action. A graphical abstract of the main novel findings is provided in Figure 8 below.

**Figure 8.**
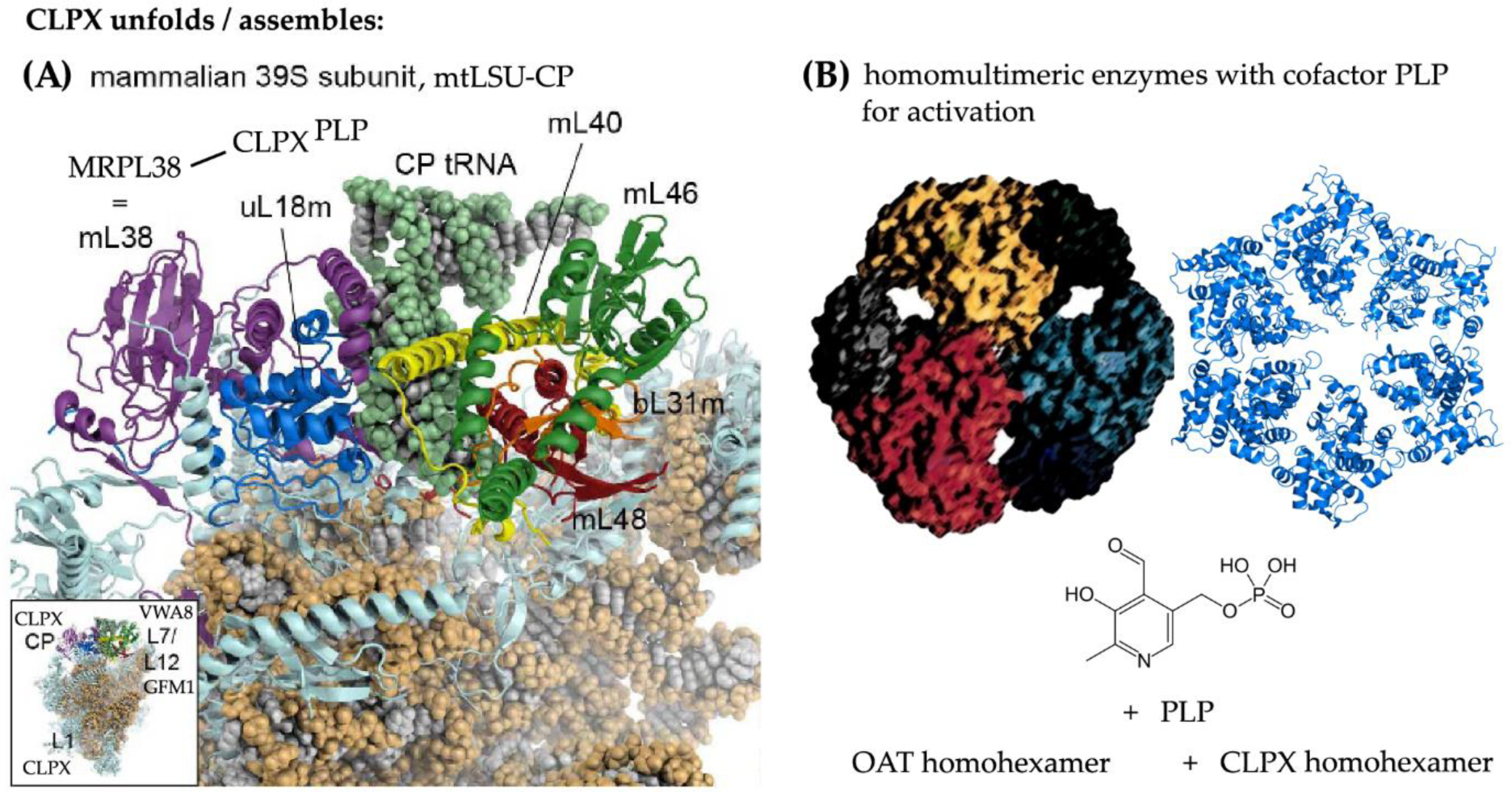
Schematic overview of endogenous CLPX (and VWA8) AAA+ unfoldase targets in the mitochondrial matrix. 3D-structure images were taken from the literature for mtLSU-CP (Extended Data Figure 8g in [79]) and for OAT homohexamer (Figure 8 in [133]), or from public databases for CLPX homohexamer (EMBL-EBI PDBe > 6sfw, last accessed on July 7, 2023) and PLP structure (EMBL-ChEBI 18405, last accessed on July 7, 2023), modifying them to compose this synopsis

### 4.1. The putative role of CLPP-dependent VWA8 for the mtLSU

The first major observation of our study (Figures 2 / 6) was the CLPP-dependent accumulation and transcriptional induction of the AAA+ unfoldase VWA8. Comigration of VWA8 was observed with mtLSU-CP intermediate assemblies. On the one hand, these include the L7/L12 stalk factors MPRL19-MRPL12-MRPL53, where GFM1 has its docking site [83] (Figure 8, inset in lower left corner). On the other hand, they include NPET wall factors MRPL23-MRPL24 with MRPL65/S30, which accumulate strongly upon CLPP depletion. Their accumulation and distortion may reflect altered peptide exit channel geometry, which may contribute to the misfolding/aggregation of mitochondrially translated proteins that characterize CLPP-mutant organelles [11, 134–136], and to the respiratory supercomplex assembly deficits observed in this study. It is important to note that the *Vwa8* transcript is induced in testis already before diplotene pathology occurs, suggesting that the VWA8 accumulation is a compensatory effort by cells in response to inadequate CLPX action. *Clpx* transcript is similarly induced at the earliest testis P17 stage as a potential reaction to functional impairment, but becomes repressed after diplotene meiosis-I failure at the P27 time point, suggesting that excess CLPX over time may acquire a toxic impact.

Very little is known about the role of VWA8 in mitochondria and peroxisomes. VWA8 contains two AAA+ domains that are relatively similar to hexameric dynein-related motors, and a VWA domain towards the C-terminus. In archaea and bacteria, each of the two domains still resides in different proteins, which interact with high frequency. AAA+ containing MoxR proteins cooperate with other VWA proteins to insert metal cofactors into substrate molecules [137–139]. In *E. coli*, this cooperation is exemplified by the concerted actions of MoxR unfoldase RavA with VWA protein ViaA, which modify cellular sensitivity to aminoglycosides antibiotics (presumably when they bind to bacterial ribosomes), or target the NADH:ubiquinone oxidoreductase-I as CI homolog, and the fumarate reductase respiratory complex [140–142]. RavA is also known for its impact on PLP-dependent lysine decarboxylase, a crucial enzyme for the generation of ornithine-derived polyamines that serve to stabilize RNA structure [143–146]. It is important to note that the closed ring RavA state bears close structural similarity to the pseudo two-fold symmetric crystal structure of bacterial ClpX [147], suggesting a common mechanism or targets. In *Acidithiobacillus ferrooxidans* bacteria the MoxR unfoldase CbbQ is upregulated upon iron exposure, and acts together with the VWA protein CbbO to rescue the CO_2_-fixing enzyme Rubisco from inactivity when its cofactor magnesium is absent [148, 149].

In a sequence analysis of AAA+ unfoldases across phylogenesis [34], the coexistence of AAA+ and VWA domains within the same protein was first observed in *Synechocystis* cyanobacteria, where the catalytic ChlD subunit of the magnesium chelatase protein complex exerts both functions. It acts to unfold tetrapyrrol ring-containing PPIX and then insert Mg^2+^ into it, thus being responsible for the first enzymatic step in chlorophyll biosynthesis [150–154]. VWA domains in prokaryotes are known only in magnesium chelatases, or in the cobaltochelatase CobT subunit, which mediates the binding of cobalt during corrin ring formation in the biosynthesis of cobalamin (also known as vitamin B12) [155–157]. Given that mammalian mitochondria do not synthesize chlorophyll or cobalamin, what related function could VWA8 fulfil to remain conserved across phylogenesis?

One possibility regarding Mg^2+^ is its decisive role for DNA/RNA conformation stability and ribosomal subunit cohesion; it controls for example also the 60-nt GTPase center of 25S rRNA in *E. coli* or the bacterial 5S rRNA loop-E stability within the LSU [158–165]. Another more relevant possibility is the fact that any metallated porphyrin system has a quadrangular shape and binds DNA/RNA avidly when four parallel strands with high guanine content adopt a G-quadruplex (G4) structure [166, 167]. G4 structures control the replication and transcription of rRNA genes, and the surface stability of both ribosome subunits [168–170]. The interaction of porphyrin variants with G4-RNA structures is among the primary pathogenesis events in some neurodegenerative diseases [171, 172]. Metallated porphyrins also have the ability to induce protein aggregation [122]. This second possibility is further supported by the further evolution of VWA functions across phylogenesis. In early eukaryotes, VWA domains become part of intracellular factors that act in DNA repair and ribosomal transport, while metazoan organisms employ VWA domains (also known as the inserted- or I-domain in integrins) always in extracellular proteins to modulate adhesion [155, 173]. The MIDAS motif (metal ion-dependent adhesion site) within the VWA domain provided the name for Midasin-1 (in yeast known as Rea1) as ancient conserved nuclear AAA-VWA protein with essential functions for cell growth, which was found to make contact with the ribosome separately via the AAA+ ring domain and via the MIDAS/VWA tail, being crucial for the rotation of the L1 stalk and the rotation of the 5S-rRNA/L5/L11 complex, to guarantee the maturation of pre-60S ribosomal LSU before nuclear export and cytosolic association with the SSU [40, 174–176].

VWA8 as mitochondrial protein that combines AAA+ and VWA domains therefore possibly functions as its prokaryotic ancestors, unfolding porphyrin or corrin rings to insert chelated magnesium or another cationic metal [177]. This notion is compatible with excess CLPX affecting heme biosynthesis, and several heavy metals accumulating with VWA8 in PRLTS3 pathology. VWA8 is localized at the matrix side of the mitochondrial inner membrane [178], and iron-chelated heme is produced inside the inner membrane in close proximity to the MICOS complex [179]. VWA8 depletion in human and zebrafish causes developmental deficits with microcephaly [180]. Heme is not only the crucial component of hemoglobin in blood cells, but also the key cofactor of cytochromes e.g. in the respiratory chain [181, 182]. Complex-IV in particular is assembled depending on the availability of heme, copper, zinc and magnesium [183–186]. This knowledge may explain the association of VWA8 with respiratory chain components (see Figure S1) and its impact there, but why would VWA8 associate selectively with the mtLSU L7/L12 stalk and NPET? The prokaryotic L7/L12 stalk is also known as the GTPase-associated center (GAC) where the translation elongation GTPases EF-G (orthologous to GFM1), IF2, RF3 and LepA dock and acquire full activity [187]. It is known that the decision whether ribosomes disassemble or alternatively intersubunit bridges get stabilized for ribosome recycling is strongly dependent on magnesium concentrations and on EF-G, which contains magnesium in its activated GTPase site [188]. At this GAC the paused translation of polyproline tracts can be rescued with the help of polyamines [145]. Regarding the association of VWA8 with the NPET, it is interesting to note that recent evidence shows the formation of G4-rRNA structures to bind heme to such an extent that this determines heme bioavailability [189, 190]. The elevated abundance of such G4 structures in CLPP-null mitoribosomes is likely, because of the observed accumulation of the G4-RNA-binding protein GRSF1 there [56]. Thus, our observation that VWA8 may interact with mtLSU proteins and with respiratory complexes is indeed compatible with current knowledge. In previous studies of mouse hepatocytes, VWA8 depletion was found to cause elevated respiratory chain complex abundance and activity, together with oxidative stress and increased protein turnover, while non-canonical gamma-amino acid GABA levels decrease [191, 192]. Thus, both experimental findings and data mining provide plausible evidence that VWA8 acts downstream from CLPX, both targeting the mitoribosome and respiratory complexes, acting probably within the homeostasis of PLP-mediated unconventional amino acids (like ALA and ornithine) and their derivatives like heme and polyamines, and the homeostasis of cationic metals, all of which are important as RNA interactors [145, 190].

### 4.2. The putative role of CLPX for the mtLSU intermediate assembly

The second major observation of our study (Figures 3 / 7) was the physiological comigration of CLPX itself with other mtLSU-CP intermediate assemblies, a completely novel finding. Upon pathological CLPP absence only specific mtLSU proteins show significant accumulation, whereas practically all mtSSU factors accumulate strongly without selectivity and without CLPX-comigration (Table S1). The selective impact of CLPXP on mtLSU was surprising, because Perrault syndrome is triggered by mutant HARS2 / LARS2 / ERAL1 all of which are associated with the mtSSU. In addition, the progressive sensorineural deafness due to the chronic administration of aminoglycoside antibiotics is also caused by neurotoxic interference and genetic variance at the mtSSU [193–195]. A detailed review of mitoribosomal pathology underlying progressive deafness as in PRLTS was recently compiled [49]. It also should be noted that the principal target of the CLPX cofactor PLP at the bacterial SSU is the inhibition of mRNA-dependent aminoacyl-tRNA binding [126]. Thus, mtSSU pathology is widespread and massive, but may not be a direct CLPXP effect, instead occurring as consequence of tRNA processing problems. The selective impact of CLPXP on mtLSU intermediate assemblies was also unexpected, because the previous concept assumed that CLPXP acts on complete ribosomes when they stall during translation. In contrast, the complexome profiles suggest that CLPX associates with specific subcomplexes either during mtLSU assembly or after mtLSU disassembly.

The CLPXP-relevant intermediate assemblies of the mtLSU were described before [72–77], and the stress-dependent redistribution of the mitochondria-specific MRPL18-L38-L40-L46-L48-L55 cluster in complexomics profiles was reported for Barth syndrome as a consequence of mitochondrial cardiolipin anomalies [73, 75, 76]. This subcomplex of the central protuberance contributes to the connection between mtLSU and mtSSU, via intersubunit bridges mB1a and mB1b from mL40-mL46-mL48 in the mtLSU to the mtSSU mitochondria-specific GTP-binding component mS29 (also abbreviated as DAP3, for Death Associated Protein 3) [196, 197]. Within this mtLSU cluster, MRPL55 (= bL31m), MRPL18 (= uL18m), MRPL38 and MRPL46 show the largest accumulation upon CLPP absence, and MRPL18 with MRPL38 exhibit most dispersion in BNE gels, in perfect overlap with CLPX dispersion.

It was unexpected that a protein cluster that did not exist in bacteria but instead arose in mitochondria should be the target of ancient CLPXP. It is therefore important to note that this MRPL18-associated protein cluster serves in bacteria as docking site for 5S rRNA but in mitochondria the cluster was modified to bind tRNA^Val^ or tRNA^Phe^ [81]. Although the peptidyltransferase activity of bacterial LSU is fulfilled by 23S rRNA in tight interaction with magnesium microclusters [198], during phylogenesis an indispensable addition occurred with 5S rRNA and its associated proteins, using allosteric effects to stabilize and coordinate various centers of the ribosome, and linking their activity with cell proliferation [199–202]. Recent observations suggest that 5S rRNA in mammalian cells can be sequestered from cytosolic ribosomes into mitochondria via interaction with proteins MRPL18 and rhodanese, where 5S rRNA can occupy its ancestral position at the mtLSU central protuberance instead of tRNA^Val/Phe^ [203–205]. Thus, the assembly of any protein with 5S-rRNA in bacteria and in mitochondria – such as MRPL18/38 - might need CLPXP.

While it is unclear what actions would be performed at this MRPL18-associated cluster, it is noteworthy that PLP as CLPX cofactor was reported to act at the bacterial LSU, dissociating the two subunits and inhibiting translation elongation, via modification of the 5S rRNA binding proteins L5-L18-L25, stalk protein L1, and elongation GTPase GFM1 ortholog [126]. Also the VWA8-comigrating ribosomal L7/L12 stalk proteins are targets of PLP modification [126]. Overall, the data raise the possibility that CLPXP with PLP as cofactor act on this mitochondria-specific cluster to control associations/disassociations of subcomplexes within the mtLSU, and of intersubunit bridges to the mtSSU. In addition, CLPXP via PLP might modulate translation efficiency via the 5S rRNA associated cluster.

The CLPX-comigrating stalk proteins MRPL1 and MRPL9 show even stronger accumulation upon CLPP deficiency, so how would they connect with such a scenario? As mentioned, L1 is a target of CLPX-cofactor PLP [126], the maturation of the L1 stalk involves a position shift during assembly steps mediated by mt-ACP2 (NDUFAB1), and the flexibility of mature L1 stalk contributes to the translocation of tRNAs during translation [80, 206]. Thus, the excess of MRPL1-L9 abundance also adds to the evidence that mutant CLPXP has an impact on translation at the elongation step.

Our demonstration that CLPX interacts with MRPL38 at endogenous abundance levels in coimmunoprecipitation assays indicates that CLPXP function for the mtLSU assembly occurs with similar abundance, as the previously observed effects of recombinant CLPX for GFM1 control in translation elongation, and as the ALAS / OAT control in the metabolism on non-classical amino acids.

### 4.3. Potential functions of CLPX and VWA8 for specific translation / RNA granule factors, matrix enzymes and PLP, as well as chaperones

A third relevant finding of our study (Figure S2 and S3) included the comigration of CLPX in WT tissue with additional specific translation factors and matrix enzymes. Among the ribonucleproteins, the histidine-tRNA synthetase HARS2, the elongation GTPase GFM1, and the cytosolic translation initiation core component EIF3C showed significant accumulation in CLPP-null tissue. HARS2 is obviously the most important result given that its mutation causes the same phenotype as CLPP deficiency, the autosomal recessive Perrault syndrome. HARS2 localizes in the mitochondrial RNA granule and associates with the mtSSU, representing a bridge between the pathology observed there and CLPXP. It is not known to associate with PLP in a specific manner, but the principal target of PLP at the bacterial SSU is the inhibition of mRNA-dependent aminoacyl-tRNA binding in general [126]. Indeed other mitochondrial amino acid-tRNA ligases also accumulate in CLPP-null tissue (in testis the increase of DARS2, EARS2, FARS2, GARS, KARS, NARS2, SARS2, TARS2, VARS2, YARS2 abundance is also >2-fold), although their physiological comigration with CLPX, and their dispersion in CLPP-null tissue are not as clear as for HARS2. Previous CLPXP target surveys throughout phylogenesis had already identified GFM1 as the most consistent result [49], and a mouse CLPXP substrate-trap screening using CLPP^WT^-FLAG versus CLPP^S149A^-FLAG plasmids transfected into CLPP-null murine embryonic fibroblasts had also identified GFM1 (named EFG1 in this report) as one of the two most credible candidates [45]. We now confirm that this interaction between CLPX and GFM1 occurs at endogenous abundance levels in coimmunoprecipitation assays even in WT tissues, and thus has physiological significance.

The mitochondrial RNA granule components PTCD1, LRPPRC, SLIRP are clearly accumulated in CLPP-null tissue, but PTCD1 was not detectable in WT complexome profiles, so its physiological CLPX-comigration remains unclear. LRPPRC migrates with SLIRP well above the physiological CLPX migration range, so their association with CLPX may occur only during pathogenesis. Thus, the RNA granule might become involved in Perrault syndrome only via off-target effects of excess CLPX. Regarding the matrix enzyme homeostasis, accumulation and CLPX-comigration of PLPBP in CLPP-null testis (severely affected in PRLTS3) but not MEF (spared in PRLTS3) (Figure S3 and Figure 5), as matrix protein responsible for the homeostasis of PLP, is apparently contributing to ALAS2 coaccumulation (see also Table S1) and increased heme production with elevated tissue iron levels. As seen in Figure 6, the mRNA for PLPBP is progressively induced during testis aging, suggesting that PLP also accumulates over time. PLP is an ancient factor that exists since the prebiotic RNA world [207], which evolved as chaperone [31] and cofactor of enzymes that mediate transaminations and other reactions such as desulfurization [208]. It is crucial for the non-classical biosynthesis of unconventional amino acids [209], e.g. when an additional amino group is placed within an amino acid at gamma-position or delta-position from the carboxy-acid group [210]. Thus, it is essential for the generation of delta-amino levulinic acid (ALA) by ALA-synthase [211], and of ornithine by ornithine-delta-aminotransferase (OAT) [212]. It is also crucial for cysteine desulfurase in the biosynthesis of molybdopterin [213], or in the FeS-cluster-mediated assembly of respiratory supercomplexes [214]. Delta-amino acids serve as building blocks in the biosynthesis of porphyrin rings (ALA), corrin rings (ALA), and polyamines (ornithine), while molybdopterin serves in the biosynthesis of the molybdenum cofactor MoCo. On these platforms heavy metals can be bound stably and harnessed for metabolic functions – porphyrin chelation of iron produces heme, corrin binding of cobalt generates cobalamin, molybdopterins stabilize molybdate. Polyamines provide heavy metal tolerance [215]. Interestingly, also the incorporation of manganese into mitochondrial superoxide dismutase was shown to depend on PLP. This occurs in a Mtm1p-dependent process in yeast, and PLP bioavailability in mitochondria depends on Mtm1p [216]. Mischarging of tRNA-Lys with ornithine occurs permanently in cells and has to be corrected by surveillance editing [217]. Its misincorporation into polypeptide chains induces a cyclic structure and translation arrest with cell toxicity [218], therefore the cleavage of such aggregation-prone fragments in mitochondria might be a function of CLPP. Jointly these lines of evidence suggest that CLPX and PLP act together in the biosynthesis of several non-classical amino acids that are needed as building blocks for the docking sites of heavy metals. Therefore, the neurodegeneration in PRLTS3 may reflect a gain-of-function of CLPX and cofactor PLP in affected tissues. Conversely, neonatal epilepsy and chronic encephalopathy due to PLPBP or PNPO loss-of-function mutations benefits from therapeutic PLP substitution [219].

In this context, it is noteworthy that also CLPX-comigrating ALDH18A1 protein multimer (also known as Glutamate-5-SemiAldehyde Synthetase, or as Pyrroline-5-Carboxylate Synthetase = P5CS, ortholog of bacterial ProA) accumulates upon CLPP absence, and was previously reported to be affected by the CLPX-cofactor PLP [220]. Proline as the most frequent cause of ribosomal stalling is metabolically derived from glutamate via Glutamate γ-SemiAldehyde (GSA) and its cyclization product Δ^1^-Pyrroline-5-Carboxylate (P5C) [221]. ALDH18A1 coaccumulates in CLPP-null tissues with Ornithine Amino Transferase (OAT) [56, 57], another proline homeostasis regulator, which is known to depend on PLP as cofactor to interconvert GSA and ornithine as intermediate enzymatic step between proline and the urea cycle [127, 128]. OAT is the mammalian ortholog of bacterial HemL (Glutamate-1-Semialdehyde-2,1-AminoMutase=GSAM), which has remarkable three-dimensional homology with ALAS, and contributes to the production of ALA with subsequent generation of tetrapyrrol rings in the heme/chlorophyll biosynthesis pathway of bacteria and plants [222–224]. OAT was proposed as CLPXP substrate in a previous study where a substrate-trap assay overexpressing CLPP^WT^-FLAG versus CLPP^S129A^-FLAG plasmids in CLPP-null MEF was combined with an N-terminomic analysis of mouse heart proteome profiles [57]. Our demonstration that CLPX interacts with OAT at endogenous abundance levels in coimmunoprecipitation assays of WT (and more strongly in CLPP-null tissue) indicates that not only ALAS association with PLP is controlled by CLPX, but a similar control mechanism is credible for OAT and its metabolic function. Our finding that *Aldh18a1* and *Oat* mRNA are induced in testis before pathology occurs, may be interpreted as cellular efforts to compensate a misassembly of the OAT homohexameric ring with PLP, with consequent impairment of its metabolic function in the interconversion of P5C and ornithine. These observations highlight similarities between proline and porphyrin biosynthesis, both starting with the conversion of GSA to cyclic pyrrol, in PLP- and CLPX-dependent pathways.

The accumulation of the molecular chaperones TRAP1 and GRP75 in the mitochondrial matrix of CLPP-null tissues may be simply due to altered assembly and slowed turnover of ribonucleoprotein complexes, as a secondary phenomenon. Furthermore, their apparent comigration with CLPX may be simply represent coincidence, given that both have a molecular weight around 70-75 kDa and their interaction with target proteins would make them migrate within the range of CLPX position dispersion. Thus, they may be purely downstream players in Perrault syndrome pathogenesis. However, it is relevant to be aware that the stress-induced cytosolic isoform of MRPL18 alters the abundance of heat shock proteins (HSP) at the level of translation, acting via its 5S rRNA association [225]. The 5S rRNA was also reported to contribute to the folding of RNA-targeting members of the HSP70 chaperone family such as DnaK in *E. coli*, the ortholog of mammalian HSPA9 in mitochondria [226]. An accumulation of the tetrapyrrol precursors of heme is also a known inducer of molecular chaperones of the HSP70 family [227]. The impact of CLPXP on MRPL18 abundance and distribution, with downstream consequences for the subcellular availability of 5S rRNA, jointly with the impact of CLPXP on the heme biosynthesis pathway, may therefore conspire to modify the stress adaptation of mitochondria and their eukaryotic hosts via molecular chaperones as an early pathogenesis event. It is important to note the high selectivity of these chaperone accumulations that constitute a pathway in themselves. As in bacteria, a conserved interaction persists between CLPX and the HSP70 family member HSPA9 with its co-chaperones GRPEL1-GRPEL2-DNLZ-CCDC127, so presumably they share specific target protein clusters.

### 4.4. Potential functions of CLPX and VWA8 for specific respiratory chain complex assembly

A fourth relevant finding concerns the question if folding problems or translation problems are primary. It is well known that (1) CLPXP targets mainly the mitoribosome, (2) CLPP mutation leads to translation fidelity deficits, (3) translation stalls mainly upon synthesis of polyproline motifs, (4) a triproline motif occurs only in mtCO1 among all mitochondrially translated factors, (5) the corresponding complex-IV activity shows the strongest reduction within the respiratory chain in CLPP-null mouse tissues. Thus, the decreased MTCO1 abundance to 30% (Figure 4) was not surprising, but a failure of assembly to respiratory supercomplexes was even stronger. Given that none of the mitochondrially translated factors combines CLPX-comigration with accumulation in the complexomics profiles (Figure S4), they are presumably affected indirectly and do not represent CLPXP targets. Two assembly factors named COX15 and SFXN4, whose accumulation and CLPX-comigration is prominent, might explain the assembly and turnover anomalies observed for complex-IV and complex-I, respectively. COX15 (also known as heme-A synthase) localizes to the inner mitochondrial membrane, where it cooperates with ferredoxin and ferredoxin reductase to convert heme-O to heme-A, which is required for the proper folding of the COX1 subunit and subsequent respiratory complex-IV assembly [112, 113, 115, 228–230]. Mutations in COX15 trigger a complex-IV deficit with subsequent infantile cardioencephalopathy that resembles Leigh syndrome [231]. Possibly explaining the complex-I deficits, SFXN4 is descended from a large family of iron transporters, impacts cytosolic iron homeostasis via the IRP1-aconitase switch, affects iron redistribution between cytosol and mitochondria, impacts mitochondrial Fe-S cluster and heme biosynthesis, assisting the MCIA complex in the assembly of the respiratory ND2 module, and influencing the levels of ferrochelatase and ALAS2 [111, 232, 233]. Its mutation triggers phenotypes of macrocytic sideroblastic anemia, weight deficits and neurological impairments [234]. Overall, the supercomplex assembly in the respiratory chain is more affected than the translation of individual proteins such as MTCO1, and the underlying protein folding problem may be reflected by the accumulation of COX15 and SFXN4 presumably due to altered homeostasis firstly of CLPX/PLP (despite compensatory VWA8 induction) and secondly of iron/heme in the mitochondrial matrix upon CLPP absence.

## 5. Conclusions

Jointly, the analyses of protein complexome profiles in mitochondria from 3 tissues in WT versus CLPP-null mice regarding accumulation, CLPX/VWA8 comigration, and dispersed gel positions have elucidated the interaction between these endogenous AAA+ unfoldases and their targets. The results highlight assembly problems at mtLSU, RNA granules, the respiratory chain complexes I and IV, and unconventional amino acid homeostasis enzymes, with consequent accumulation of molecular chaperone proteins. Practically all protein accumulations appeared to represent secondary compensatory events presumably due to assembly and function deficits, only the CLPX protein accumulation led to a compensatory transcriptional downregulation and may therefore by itself explain PRLTS3 pathology through its toxic gain-of-function. The data provide proof-of-principle that CLPX controls not only the access of PLP to ALAS2 to enhance its activity, but similarly OAT, GFM1 and MRPL38 may be modulated in this way. This provides mechanistic detail in the understanding of PRLTS3 pathogenesis where CLPX is in excess, and progressive deafness is due to mitoribosomal translation problems.

## Abbreviations

12S rRNA: ribosomal RNA component of SSU in bacteria
16S rRNA: ribosomal RNA component of LSU in bacteria
16S rRNA: ribosomal RNA component of mtSSU in mammals
23S rRNA: ribosomal RNA component of mtLSU body
28S: mtSSU of mammalian mitoribosomes
39S: mtLSU of mammalian mitoribosomes
55S: completely assembled mitoribosome
5S rRNA: ribosomal RNA component of the mtLSU central protuberance
AAA+: protein superfamily of ring-shaped P-loop NTPases
ABC: AmmoniumBiCarbonate
ACAD9: Acyl-Coenzyme A Dehydrogenase Family, Member 9, MCIA component
ACN: Acetonitrile
ACTB: beta-Actin protein
AFG3L2: ATPase Family Member 3-Like 2, Matrix AAA Peptidase Subunit 2
AGC: Automatic gain control
ALA: delta-Amino Levulinic Acid
ALAS: 5-AminoLevulinic Acid Synthase
ALAS2: Delta-AminoLevulinic-Acid-Synthase-2, erythroid-specific, mitoch.
ALDH18A1: =P5CS, Aldehyde Dehydrogenase 18 Family Member A1
ATAD3: ATPase Family AAA Domain-Containing Protein 3
ATPase: Adenosine 5’-TriPhosphatase
BCA: BiCinchoninic Acid
BCS1L: Ubiquinol-Cytochrome C Reductase Complex Chaperone
BNE: Blue Native gel Electrophoresis
CaCl_2_: Calcium chloride
CbbO: Prokaryotic Rubisco activation protein
CbbQ: Prokaryotic Nitric oxide reductase NorQ protein; AAA ATPase
cDNA: complementary DeoxyriboNucleic Acid
ChlD: Prokaryotic Magnesium protoporphyrin IX chelatase, d subunit
CI: Respiratory Complex-I
CII: Respiratory Complex-II
CIII: Respiratory Complex-III
CIV: Respiratory Complex-IV
CCDC127: Coiled-Coil Domain-Containing Protein 127
ClpA: Prokaryotic ATP-dependent specificity component of ClpAP protease
CLPB: Caseinolytic peptidase B protein homolog; AAA+ ATPase
CLPP: ATP-dependent Clp protease Proteolytic subunit, mitochondrial
CLPX: specificity component of Clp protease complex, AAA+ ATPase
CLPXP: proteolytic machine with components CLPP and CLPX
CO_2_: Carbon dioxide
CobT: Prokaryotic Cobaltochelatase
CoIP: Co-immunoprecipitation
*Cox1*: Cytochrome-c-OXidase subunit 1, catalytic
COX15: Cytochrome-c-OXidase assembly-factor; acts in heme-A biosynthesis
CP: Central Protuberance of mtLSU
CV: Respiratory Complex-V
DNA: DeoxyriboNucleic Acid
DnaJ: Prokaryotic co-chaperone Hsp40; acts with DnaK/GrpE in disassembly
DnaK: Prokaryotic chaperone Hsp70
DAP3: =MRPS29, Death Associated Protein 3
DNLZ: DNL-type Zinc finger Protein; mtHsp70-escort protein
DTT: DiThioThreitol
ECSIT: Evolutionary Conserved Signal Intermediate In Toll pathway, mitoch.
EDTA: EthyleneDiamineTetraacetic Acid
EF-G: Translation Elongation Factor G, ortholog of GFM1
EIF3: Eukaryotic translation Initiation Factor 3 protein complex
EIF3C: Eukaryotic translation Initiation Factor 3, subunit C
ERAL1: Era-Like 12S mitochondrial rRNA chaperone 1
FASTKD4: =TBRG4, FAST Kinase Domain-Containing Protein 4
FC: Fold-Change
FDR: False Discovery Rate
FECH: FErroCHelatase, inserting iron into porphyrin rings to produce heme
Fe-S cluster: Iron-Sulfur cluster
FLAG: Octapeptide tag for proteins, with sequence DYKDDDDK
G4: Guanosine-rich quadruplex structure of DNA/RNA
GABA: Gamma-AminoButyric Acid, main inhibitory neurotransmitter
GAC: GTPase-AssociatedC
GAPDH: GlycerAldehyde 3-Phosphate DeHydrogenase, glycolysis enzyme
GFM1: translation elongation Factor G 1, Mitochondrial
GFM2: translation elongation Factor G 2, Mitochondrial
GRPEL1: GrpE-Like 1 protein co-chaperone, mitochondrial
GRPEL2: GrpE-Like 2 protein co-chaperone, mitochondrial
GRSF1: G-Rich RNA Sequence binding Factor 1
GroEL: Prokaryotic chaperonin Hsp60, peptide-dependent ATPase
GSA: Glutamate γ-SemiAldehyde
GSAM: Glutamate-1-SemiAldehyde-2,1-AminoMutase
GTP: Guanosine 5’-TriPhosphate
GTPase: Guanosine 5’-TriPhosphatase
HARS2: Histidine-tRNA synthase, mitochondrial
HemL: Prokaryotic ortholog of GSAM, heme/porphyrin biosynthesis enzyme
HPLC: High-Performance Liquid Chromatography
Hsp100: Eukaryotic Heat shock protein family, ∼100 kDa size in yeast
HSP60: Chaperonin family of Heat Shock Proteins, ∼60 kDa size in bacteria
Hsp70: Ubiquitous family of Heat shock proteins, homologous to DnaK
HSP90: Bacterial/eukaryotic family of Heat Shock Proteins
HSPA9: =Mortalin/GRP75, Heat Shock Protein family A (Hsp70) member 9, mito.
HssR: Prokaryotic Heme response Regulator
IBAQ: Intensity-Based Absolute Quantification
ICP-MS: Inductively coupled plasma mass spectrometry
ID: IDentity code
IF2: Prokaryotic translation Initiation Factor 2, protects formylMet-tRNA
InfB: Prokaryotic translation INitiation Factor if-2
IRP1: =IREB1/Aconitase-1, Iron Regulatory Protein 1
kDa: kiloDalton
KGD4: =MRPS36, Alpha-Ketoglutarate Dehydrogenase Component 4
LARS2: Leucine-tRNA synthase, mitochondrial
LC/MS: Liquid Chromatography / Mass Spectrometry
LepA: Prokaryotic elongation factor, back-translocating, stress fidelity, GTPase
LONP1: Lon Peptidase 1, Mitochondrial
LRPPRC: Leucine Rich Pentatricopeptide Repeat Containing
MCIA: Mitochondrial Complex-I Assembly complex
MDa: MegaDalton
MEF: Murine Embryonic Fibroblasts
MICOS: MItochondrial Contact site and cristae Organizing System
MIDAS: Metal Ion-Dependent Adhesion Site
Mg^2+^: Magnesium ion
mM: milliMolar
MoxR: Prokaryotic magnesium chelatase, AAA+ ATPase
MRPL18: Mitochondrial Ribosomal Protein L18
MRPL38: Mitochondrial Ribosomal Protein L38
MRPP1: =TRMT10C, Mitochondrial Ribonuclease P Protein 1
MS: Mass Spectrometry
MSMS: tandem Mass Spectrometry
MT-ATP8: Mitochondrially encoded ATP synthase membrane subunit 8
MTCO1: =COX1, Mitochondrially encoded Cytochrome c Oxidase I
MTCO2: =COX2, Mitochondrially encoded Cytochrome c Oxidase II
MTCO3: =COX3, Mitochondrially encoded Cytochrome c Oxidase III
MTHFD2: MethyleneTetraHydroFolate Dehydrogenase (NADP+ dependent) 2
mtLSU: mitochondrial Large SubUnit of ribosome
MTND3: Mitochondrially encoded NADH Dehydrogenase 3
MTND4: Mitochondrially encoded NADH Dehydrogenase 4
MTND5: Mitochondrially encoded NADH Dehydrogenase 5
MTND6: Mitochondrially encoded NADH Dehydrogenase 6
mtSSU: mitochondrial Small SubUnit of ribosome
m/z: ratio of mass versus charge number of ions
NaCl: Sodium Chloride salt
NADH: reduced form of Nicotinamide Adenine Dinucleotide
NDUFAB1: =mt-ACP2, NADH:Ubiquinone oxidoreductase subunit AB1
NDUFAF1: NADH:Ubiquinone oxidoreductase complex Assembly Factor 1
NDUFAF3: NADH:Ubiquinone oxidoreductase complex Assembly Factor 3
NDUFS1: NADH:Ubiquinone oxidoreductase core subunit S1
NDUFV1: NADH:Ubiquinone oxidoreductase core subunit V1
NDUFV2: NADH:Ubiquinone oxidoreductase core subunit V2
NP40: Nonyl Phenoxypolyethoxylethanol, non-ionic, non-denaturing detergent
NPET: Nascent Polypeptide Exit Tunnel in large subunit of ribosome
NTPase: Nucleoside-TriPhosphatase
OAT: Ornithine delta-AminoTransferase, mitochondrial
p: Probability of error
P21: Postnatal day 21
P5C: Pyrroline-5-Carboxylate
P5CS: =ALDH18A1, Pyrroline-5-Carboxylate Synthetase
PARL: Presenilin-Associated-Rhomboid-Like, mito. intramembrane protease
PBS: Phosphate-Buffered Saline solution
PBS/T: Phosphate-Buffered Saline solution with Tween-20
PCR: Polymerase Chain Reaction
PDK1: Pyruvate Dehydrogenase Kinase 1
PDK3: Pyruvate Dehydrogenase Kinase 3
PDPR: Pyruvate Dehydrogenase Phosphatase Regulatory subunit
pH: “potential of Hydrogen”, scale to quantify acidity of aqueous solutions
PHB1: ProHiBitin-1, forms large ring complexes in mito. inner membrane
PLP: =P5P, PyridoxaL 5’-Phosphate, vitamin B6 active form, enzyme cofactor
PLPBP: =PROSC, PyridoxaL Phosphate Binding Protein
POLDIP2: DNA Polymerase Delta Interacting Protein 2
PP/PE: PolyPropylen/PolyEthylene
PPIX: ProtoPorphyrin IX
PPOX: ProtoPorphyrin IX oxidase
PRIDE: Proteomics Identification Database – EMBL/EBI
PRLTS3: PerRauLT Syndrome type 3
ProA: Prokaryotic ortholog of ALDH18A1/P5CS
PRORP: =MRPP3, Protein Only RNase-P Catalytic Subunit
PTCD1: PentaTriCopeptide repeat Domain 1
PTCD2: PentaTriCopeptide repeat Domain 2
PTCD3: =MRPS39/COXPD51, PentaTriCopeptide repeat Domain 3
RavA: Prokaryotic AAA+ moxr family ATPase, putative molecular chaperone
Rea1: =Midasin-1, yeast chaperone for maturation of pre-60S ribosome
RF3: Prokaryotic translation Release Ractor 3
RIPA: Radio-Immuno-Precipitation Assay buffer
RMND1: Required for Meiotic Nuclear Division 1 homolog
RNA: Ribo-Nucleic Acid
RpsA: Prokaryotic RNA chaperone to unfold structured mRNA on ribosome
rRNA: ribosomal RNA
RT: Room Temperature
RT-qPCR: Reverse Transcriptase real-time quantitative PCR
SC: SuperComplex
SDS: Sodium Dodecyl Sulfate, detergent
SEM: Standard Error of the Mean
SFXN4: SideroFleXiN 4
SHMT2: Serine HydroxyMethylTransferase 2
SLC25A28: SoLute Carrier family 25 member 28, mitochondrial iron transporter
SLIRP: SRA Stem-Loop Interacting RNA-binding Protein, mitochondrial
SLP2: =STOML2, Stomatin-Like Protein 2, at mitoch. inner membrane rafts
SPATA5: Spermatogenesis-associated factor protein, mito. ribosome maturation
SPG7: Matrix-AAA peptidase subunit, paraplegin
STRING: Search Tool for the Retrieval of INteracting genes/Proteins
T: statistical Trend
*Tbp*: TATA-Binding Protein
TBRG4: =FASTKD4, Transforming growth factor Beta ReGulator 4
TIMMDC1: Translocase of Inner Mitochondrial Membrane Domain Containing 1
TMEM126B: TransMEMbrane protein 126B, mitochondrial complex-I assembly factor
TMEM186: TransMEMbrane protein 186, mitochondrial complex-I assembly factor
TRAP1: =HSP90L, TNF Receptor Associated Protein 1, mitochondrial chaperone
TRMT10C: =MRPP1, tRNA Methyltransferase 10C, mitochondrial RNase-P Subunit
tRNA: transfer RNA
tRNA^Phe^: tRNA for the amino acid Phenylalanine
tRNA^Val^: tRNA for the amino acid Valine
TUBA4A: TUBulin Alpha 4a protein
TWNK: TWiNKle mtDNA helicase, Ataxin-8
UPR: Unfolded Protein Response pathway
UPRmt: Unfolded Protein Response in mitochondria
v/v: ratio volume per volume
VCP: Valosin Containing Protein, transitional ER AAA+ ATPase
ViaA: Prokaryotic VWA-domain-protein, CoxE-like family, AAA+ interacting
VWA: von Willebrand factor type A domain
VWA8: Von Willebrand factor A domain containing 8
WT: wildtype
w/v: ratio weight per volume
YME1L1: ATP-dependent zinc metalloprotease, mitoch. intermembrane space
ZFE: Central Animal Facility, University Hospital Frankfurt am Main

## Acknowledgments

We thank Heike Angerer for technical support regarding the purification of mitochondria, Hildegard König regarding our biosafety S1/S2 lab, and the staff of the animal facility ZFE in Frankfurt/M. We are very grateful to Dr. Ilka Wittig and Jana Meisterknecht in the Functional Proteomics team at Goethe University for excellent mass spectrometry services. We thank Michael Gertitschke and Andrea Reichlmeir for great laboratory assistance.

## Data Availability

All primary data were made publically available by the ProteomeXchange Consortium via the PRIDE partner repository with the dataset identifiers PXD035352 (testis), PXD036901 (brain) and PXD036933 (heart).

## Author Contributions

Conceptualization, J.K. and G.A.; methodology, J.K., S.G., G.K., J.S.-W., and M.R.,; software, J.K. and G.A.; validation, J.K., S.G., G.K., J.S.-W., M.R., and G.A.; formal analysis, J.K., J.S.-W. and G.A.; investigation, J.K., S.G., J.S.-W. and G.A.; resources, J.S.-W. and G.A.; data curation, J.K.; writing—original draft preparation, J.K., J.S.-W. and G.A.; writing—review and editing, J.K., J.S.-W. and G.A.; visualization, J.K. and G.A.; supervision, J.K., S.G., J.S.-W. and G.A.; project administration, G.A.; funding acquisition, J.S.-W. and G.A. All authors have read and agreed to the published version of the manuscript.

## Funding

The metal analyses in mouse brains were funded by the core budget of JSW.

## Institutional Review Board Statement

The study was conducted according to the guidelines of the Declaration of Helsinki, and approved by the Institutional Review Board of Regierungspräsidium Darmstadt (protocol code V54-19c20/15-FK/1073, date of approval Sept. 28, 2016).

## Conflicts of Interest

The authors declare no conflict of interest. The funders had no role in the design of the study; in the collection, analyses, or interpretation of data; in the writing of the manuscript, or in the decision to publish the results.

**Supplementary Materials: Figure S1:** VWA8 as prey in human protein-protein-interactions, **Figure S2:** Complexomics profiles of ribonucleoproteins, **Figure S3:** Complexomics profiles of mitochondrial matrix PLP-associated factors and other chaperones, **Figure S4:** Complexomics profiles of the detected mitochondrially translated OXPHOS complex components and relevant assembly factors, **Table S1:** Complexomics profiles from testis, brain and heart mitochondria. **Table S2:** Heavy metal levels in mouse brain tissue.

**Figure S1. VWA8 as prey in human protein-protein-interactions**, according to BIOGRID database, visualized and analyzed for pathway enrichments by the STRING webserver. To avoid overexpression artifacts where excess VWA8 remains excluded from mitochondrial import and interacts with cytosolic or nuclear factors, only the BIOGRID dataset on VWA8 as prey was taken for analysis, not the dataset with VWA8 as bait. Practically all interactors are known for their localization to the mitochondrial matrix or inner membrane. Their overall protein-protein-interaction enrichment is highly significant (<1.0e-16), as illustrated by lines that highlight interactions previously known to STRING in several colors (each color representing different technical approaches of ascertainment and credibility). The interacting proteins are illustrated as buttons where different colors reflect their involvement in assembly complexes (green color) or pathways, e.g. the mitoribosomal translation apparatus (red), the respiratory chain complexes I-V (blue), RNA-binding proteins many of which cluster in the mitochondrial RNA granule (yellow). The significance of enrichment is analyzed in lines at the lower figure end. In columns, the webserver shows for each Gene Ontology (GO) term (1) the number, (2) the description, (3) the count of factors observed among all factors in the network, (4) the enrichment strength, (5) the false discovery rate (FDR). As shown, the BIOGRID database comprises VWA8 interaction data well beyond previous STRING interaction knowledge, and indicates the association of the VWA8 unfoldase with the mitochondrial RNA granule, the mitoribosomal translation machinery, several respiratory chain complexes, and various metabolic enzymes.

**Figure S2. Complexomics profiles of ribonucleoproteins** from the mitochondrial matrix that showed CLPX/VWA8 comigration/dispersion as well as accumulated abundance in CLPP-null tissues. The format is analogous to Figure 2. Overall, the physiological CLPX and VWA8 migration range (dark grey box) overlaps with several mRNA translation factors (GFM1, EIF3C, HARS2, TARS2), while the dispersed CLPX (light grey box) and VWA8 migration range also includes various RNA processing factors (LRPPRC, SLIRP, MTHFD2, PTCD1, PTCD2, TRMT10C/MRPP1, GRSF1, FASTKD2, PDE12).

**Figure S3. Complexomics profiles of mitochondrial matrix PLP-associated factors and other chaperones** that showed CLPX/VWA8 comigration/dispersion as well as accumulated abundance in CLPP-null tissues. The format is analogous to Figure 2. Overall, the dispersed CLPX (light grey box) migration range overlaps with PLP-associated proteins (PLPBP, ALAS2, OAT) and with several other chaperone-pathway factors, but also with unrelated pyruvate-homeostasis enzymes.

**Figure S4.** Complexomics profiles of the detected mitochondrially translated OXPHOS complex components and relevant assembly factors, regarding CLPX/VWA8 comigration/dispersion as well as accumulated abundance in CLPP-null tissues. Three matrix-immersed complex-I N-module core factors (S1, V1 and V2) that were shown to depend on CLPP in their assem-bly/disassembly, and the 12 detected factors that are mitochondrially encoded/translated within respiratory chain complexes I/III/IV/V were studied. Fold changes were calculated (1) across all slices per tissue (total amount), (2) as average of tissues, (3) in the CLPX migration range from slice 4 to 28, (4) in the fully assembled complex from slice 29 to 48. The color code is analogous to Figure 2. Comigration with CLPX was detected mainly for the N-module factors and for MTCO1-3 (COX1-3), but none of them showed accumulation upon CLPP deficiency. The N-module factors accumulated selectively within the CLPX comigration range, while MTCO1-3 were reduced also there. Integration in fully assembled complexes was reduced for N-module factors and for almost all mitochondrially encoded proteins, reaching values of almost 0 for MTCO1-3. Among complex-IV components, COX15 was conspicuous for its comigration with CLPX and its strong accumulation in CLPP-null tissues. When individual factors due to scarce abundance were not detectable at all by mass spectrometry in a WT sample, the resulting FC values of infinite upregulation in mutant can be considered artefacts.

**Table S1:** Complexomics profiles from testis, brain and heart mitochondria.

**Table S2:** Heavy metal levels in mouse brain tissue.

## Notes

### Competing Interest Statement

The authors have declared no competing interest.

## References

1. Schirmer, E.C., et al., HSP100/Clp proteins: a common mechanism explains diverse functions. Trends Biochem Sci, 1996. 21(8): p. 289–96.

2. Olivares, A.O., T.A. Baker, and R.T. Sauer, Mechanical Protein Unfolding and Degradation. Annu Rev Physiol, 2018. 80: p. 413–429.

3. De Gaetano, A., et al., Impaired Mitochondrial Morphology and Functionality in Lonp1(wt/-) Mice. J Clin Med, 2020. 9(6).

4. Bezawork-Geleta, A., et al., LON is the master protease that protects against protein aggregation in human mitochondria through direct degradation of misfolded proteins. Sci Rep, 2015. 5: p. 17397.

5. Shin, M., et al., Structures of the human LONP1 protease reveal regulatory steps involved in protease activation. Nat Commun, 2021. 12(1): p. 3239.

6. Park, S.C., et al., Oligomeric structure of the ATP-dependent protease La (Lon) of Escherichia coli. Mol Cells, 2006. 21(1): p. 129–34.

7. Fischer, F., et al., Human CLPP reverts the longevity phenotype of a fungal ClpP deletion strain. Nat Commun, 2013. 4: p. 1397.

8. Haynes, C.M., et al., ClpP mediates activation of a mitochondrial unfolded protein response in C. elegans. Dev Cell, 2007. 13(4): p. 467–80.

9. Zhao, Q., et al., A mitochondrial specific stress response in mammalian cells. EMBO J, 2002. 21(17): p. 4411–9.

10. Gersch, M., et al., AAA+ chaperones and acyldepsipeptides activate the ClpP protease via conformational control. Nat Commun, 2015. 6: p. 6320.

11. Wawrzynow, A., et al., The ClpX heat-shock protein of Escherichia coli, the ATP-dependent substrate specificity component of the ClpP-ClpX protease, is a novel molecular chaperone. EMBO J, 1995. 14(9): p. 1867–77.

12. Laut, C.L., et al., DnaJ and ClpX Are Required for HitRS and HssRS Two-Component System Signaling in Bacillus anthracis. Infect Immun, 2022. 90(1): p. e0056021.

13. Mangla, N., R. Singh, and N. Agarwal, HtpG Is a Metal-Dependent Chaperone Which Assists the DnaK/DnaJ/GrpE Chaperone System of Mycobacterium tuberculosis via Direct Association with DnaJ2. Microbiol Spectr, 2023. 11(3): p. e0031223.

14. Farrand, A.J., et al., Regulation of host hemoglobin binding by the Staphylococcus aureus Clp proteolytic system. J Bacteriol, 2013. 195(22): p. 5041–50.

15. Farrand, A.J., et al., Proteomic analyses of iron-responsive, Clp-dependent changes in Staphylococcus aureus. Pathog Dis, 2015. 73(3).

16. Kobe, C., et al., Perrault Syndrome with progressive nervous system involvement. Clin Nucl Med, 2008. 33(12): p. 922–4.

17. Key, J., et al., CLPP Depletion Causes Diplotene Arrest; Underlying Testis Mitochondrial Dysfunction Occurs with Accumulation of Perrault Proteins ERAL1, PEO1, and HARS2. Cells, 2022. 12(1).

18. Faridi, R., et al., New insights into Perrault syndrome, a clinically and genetically heterogeneous disorder. Hum Genet, 2022. 141(3-4): p. 805–819.

19. Hochberg, I., et al., Bi-allelic variants in the mitochondrial RNase P subunit PRORP cause mitochondrial tRNA processing defects and pleiotropic multisystem presentations. Am J Hum Genet, 2021. 108(11): p. 2195–2204.

20. Yien, Y.Y., et al., Mutation in human CLPX elevates levels of delta-aminolevulinate synthase and protoporphyrin IX to promote erythropoietic protoporphyria. Proc Natl Acad Sci U S A, 2017. 114(38): p. E8045–E8052.

21. Ducamp, S., et al., A mutation in the iron-responsive element of ALAS2 is a modifier of disease severity in a patient suffering from CLPX associated erythropoietic protoporphyria. Haematologica, 2021. 106(7): p. 2030–2033.

22. van der Vorm, L.N. and B.H. Paw, Studying disorders of vertebrate iron and heme metabolism using zebrafish. Methods Cell Biol, 2017. 138: p. 193–220.

23. Shanley, B.C., et al., Neurochemical aspects of porphyria. Studies on the possible neurotoxicity of delta-aminolaevulinic acid. S Afr Med J, 1975. 49(14): p. 576–80.

24. Jimenez-Jimenez, F.J., et al., Hereditary Coproporphyria Associated with the Q306X Mutation in the Coproporphyrin Oxidase Gene Presenting with Acute Ataxia. Tremor Other Hyperkinet Mov (N Y), 2013. 3.

25. Kevelam, S.H., et al., Acute intermittent porphyria-related leukoencephalopathy. Neurology, 2016. 87(12): p. 1258–65.

26. Yasuda, M., et al., Homozygous hydroxymethylbilane synthase knock-in mice provide pathogenic insights into the severe neurological impairments present in human homozygous dominant acute intermittent porphyria. Hum Mol Genet, 2019. 28(11): p. 1755–1767.

27. Ferreira, G.C., et al., Iron Hack - A symposium/hackathon focused on porphyrias, Friedreich’s ataxia, and other rare iron-related diseases. F1000Res, 2019. 8: p. 1135.

28. Stutterd, C.A., et al., Expanding the clinical and radiological phenotypes of leukoencephalopathy due to biallelic HMBS mutations. Am J Med Genet A, 2021. 185(10): p. 2941–2950.

29. Sedel, F., [Inborn errors of metabolism in adult neurology]. Rev Neurol (Paris), 2013. 169 Suppl 1: p. S63–9.

30. Kardon, J.R., et al., Mitochondrial ClpX Activates a Key Enzyme for Heme Biosynthesis and Erythropoiesis. Cell, 2015. 161(4): p. 858–67.

31. Cellini, B., et al., The chaperone role of the pyridoxal 5’-phosphate and its implications for rare diseases involving B6-dependent enzymes. Clin Biochem, 2014. 47(3): p. 158–65.

32. Furuyama, K., K. Kaneko, and P.D. Vargas, Heme as a magnificent molecule with multiple missions: heme determines its own fate and governs cellular homeostasis. Tohoku J Exp Med, 2007. 213(1): p. 1–16.

33. Hedtke, B., et al., HEMA RNAi silencing reveals a control mechanism of ALA biosynthesis on Mg chelatase and Fe chelatase. Plant Mol Biol, 2007. 64(6): p. 733–42.

34. Neuwald, A.F., et al., AAA+: A class of chaperone-like ATPases associated with the assembly, operation, and disassembly of protein complexes. Genome Res, 1999. 9(1): p. 27–43.

35. Mabanglo, M.F. and W.A. Houry, Recent structural insights into the mechanism of ClpP protease regulation by AAA+ chaperones and small molecules. J Biol Chem, 2022. 298(5): p. 101781.

36. Kim, Y.I., et al., Molecular determinants of complex formation between Clp/Hsp100 ATPases and the ClpP peptidase. Nat Struct Biol, 2001. 8(3): p. 230–3.

37. Kang, S.G., et al., Human mitochondrial ClpP is a stable heptamer that assembles into a tetradecamer in the presence of ClpX. J Biol Chem, 2005. 280(42): p. 35424–32.

38. Gribun, A., et al., The ClpP double ring tetradecameric protease exhibits plastic ring-ring interactions, and the N termini of its subunits form flexible loops that are essential for ClpXP and ClpAP complex formation. J Biol Chem, 2005. 280(16): p. 16185–96.

39. Baker, T.A. and R.T. Sauer, ClpXP, an ATP-powered unfolding and protein-degradation machine. Biochim Biophys Acta, 2012. 1823(1): p. 15–28.

40. Prattes, M., et al., Shaping the Nascent Ribosome: AAA-ATPases in Eukaryotic Ribosome Biogenesis. Biomolecules, 2019. 9(11).

41. de Lonlay, P., et al., A mutant mitochondrial respiratory chain assembly protein causes complex III deficiency in patients with tubulopathy, encephalopathy and liver failure. Nat Genet, 2001. 29(1): p. 57–60.

42. Juhola, M.K., et al., The mitochondrial inner membrane AAA metalloprotease family in metazoans. FEBS Lett, 2000. 481(2): p. 91–5.

43. Escobar-Henriques, M. and V. Anton, Mitochondrial Surveillance by Cdc48/p97: MAD vs. Membrane Fusion. Int J Mol Sci, 2020. 21(18).

44. Issop, L., et al., Mitochondria-associated membrane formation in hormone-stimulated Leydig cell steroidogenesis: role of ATAD3. Endocrinology, 2015. 156(1): p. 334–45.

45. Szczepanowska, K., et al., CLPP coordinates mitoribosomal assembly through the regulation of ERAL1 levels. EMBO J, 2016. 35(23): p. 2566–2583.

46. Dennerlein, S., et al., Human ERAL1 is a mitochondrial RNA chaperone involved in the assembly of the 28S small mitochondrial ribosomal subunit. Biochem J, 2010. 430(3): p. 551–8.

47. Uchiumi, T., et al., ERAL1 is associated with mitochondrial ribosome and elimination of ERAL1 leads to mitochondrial dysfunction and growth retardation. Nucleic Acids Res, 2010. 38(16): p. 5554–68.

48. Key, J., et al., Global Proteome of LonP1(+/-) Mouse Embryonal Fibroblasts Reveals Impact on Respiratory Chain, but No Interdependence between Eral1 and Mitoribosomes. Int J Mol Sci, 2019. 20(18).

49. Auburger, G., J. Key, and S. Gispert, The Bacterial ClpXP-ClpB Family Is Enriched with RNA-Binding Protein Complexes. Cells, 2022. 11(15).

50. Rumyantseva, A., M. Popovic, and A. Trifunovic, CLPP deficiency ameliorates neurodegeneration caused by impaired mitochondrial protein synthesis. Brain, 2022. 145(1): p. 92–104.

51. Gispert, S., et al., Loss of mitochondrial peptidase Clpp leads to infertility, hearing loss plus growth retardation via accumulation of CLPX, mtDNA and inflammatory factors. Hum Mol Genet, 2013. 22(24): p. 4871–87.

52. Lee, M., et al., Reconstitution of mammalian mitochondrial translation system capable of correct initiation and long polypeptide synthesis from leaderless mRNA. Nucleic Acids Res, 2021. 49(1): p. 371–382.

53. Szczepanowska, K., et al., A salvage pathway maintains highly functional respiratory complex I. Nat Commun, 2020. 11(1): p. 1643.

54. Mabanglo, M.F., V. Bhandari, and W.A. Houry, Substrates and interactors of the ClpP protease in the mitochondria. Curr Opin Chem Biol, 2022. 66: p. 102078.

55. Bhandari, V., et al., The Role of ClpP Protease in Bacterial Pathogenesis and Human Diseases. ACS Chem Biol, 2018. 13(6): p. 1413–1425.

56. Key, J., et al., Inactivity of Peptidase ClpP Causes Primary Accumulation of Mitochondrial Disaggregase ClpX with Its Interacting Nucleoid Proteins, and of mtDNA. Cells, 2021. 10(12).

57. Hofsetz, E., et al., The Mouse Heart Mitochondria N Terminome Provides Insights into ClpXP-Mediated Proteolysis. Mol Cell Proteomics, 2020. 19(8): p. 1330–1345.

58. Strack, P.R., et al., Polymerase delta-interacting protein 38 (PDIP38) modulates the stability and activity of the mitochondrial AAA+ protease CLPXP. Commun Biol, 2020. 3(1): p. 646.

59. Lasserre, J.P., et al., A complexomic study of Escherichia coli using two-dimensional blue native/SDS polyacrylamide gel electrophoresis. Electrophoresis, 2006. 27(16): p. 3306–21.

60. Wittig, I., H.P. Braun, and H. Schagger, Blue native PAGE. Nat Protoc, 2006. 1(1): p. 418–28.

61. Cox, J. and M. Mann, MaxQuant enables high peptide identification rates, individualized p.p.b.-range mass accuracies and proteome-wide protein quantification. Nat Biotechnol, 2008. 26(12): p. 1367–72.

62. Perez-Riverol, Y., et al., The PRIDE database resources in 2022: a hub for mass spectrometry-based proteomics evidences. Nucleic Acids Res, 2022. 50(D1): p. D543–D552.

63. Giese, H., et al., NOVA: a software to analyze complexome profiling data. Bioinformatics, 2015. 31(3): p. 440–1.

64. Schlattl, M., M. Buffler, and W. Windisch, Clay Minerals Affect the Solubility of Zn and Other Bivalent Cations in the Digestive Tract of Ruminants In Vitro. Animals (Basel), 2021. 11(3).

65. Adrain, C., E.M. Creagh, and S.J. Martin, Apoptosis-associated release of Smac/DIABLO from mitochondria requires active caspases and is blocked by Bcl-2. EMBO J, 2001. 20(23): p. 6627–36.

66. Baghirova, S., et al., Sequential fractionation and isolation of subcellular proteins from tissue or cultured cells. MethodsX, 2015. 2: p. 440–5.

67. Livak, K.J. and T.D. Schmittgen, Analysis of relative gene expression data using real-time quantitative PCR and the 2(-Delta Delta C(T)) Method. Methods, 2001. 25(4): p. 402–8.

68. Maurizi, M.R., et al., Endopeptidase Clp: ATP-dependent Clp protease from Escherichia coli. Methods Enzymol, 1994. 244: p. 314–31.

69. Wai, T., et al., The membrane scaffold SLP2 anchors a proteolytic hub in mitochondria containing PARL and the i-AAA protease YME1L. EMBO Rep, 2016. 17(12): p. 1844–1856.

70. Liu, H., et al., Prohibitin 1 regulates mtDNA release and downstream inflammatory responses. EMBO J, 2022. 41(24): p. e111173.

71. Manon, S., M. Priault, and N. Camougrand, Mitochondrial AAA-type protease Yme1p is involved in Bax effects on cytochrome c oxidase. Biochem Biophys Res Commun, 2001. 289(5): p. 1314–9.

72. Wessels, H.J., et al., Analysis of 953 human proteins from a mitochondrial HEK293 fraction by complexome profiling. PLoS One, 2013. 8(7): p. e68340.

73. Gardeitchik, T., et al., Bi-allelic Mutations in the Mitochondrial Ribosomal Protein MRPS2 Cause Sensorineural Hearing Loss, Hypoglycemia, and Multiple OXPHOS Complex Deficiencies. Am J Hum Genet, 2018. 102(4): p. 685–695.

74. Chatzispyrou, I.A., et al., A homozygous missense mutation in ERAL1, encoding a mitochondrial rRNA chaperone, causes Perrault syndrome. Hum Mol Genet, 2017. 26(13): p. 2541–2550.

75. Van Strien, J., et al., COmplexome Profiling ALignment (COPAL) reveals remodeling of mitochondrial protein complexes in Barth syndrome. Bioinformatics, 2019. 35(17): p. 3083–3091.

76. Aibara, S., et al., Structural basis of mitochondrial translation. Elife, 2020. 9.

77. Bogenhagen, D.F., et al., Kinetics and Mechanism of Mammalian Mitochondrial Ribosome Assembly. Cell Rep, 2018. 22(7): p. 1935–1944.

78. Brown, A., et al., Structure of the large ribosomal subunit from human mitochondria. Science, 2014. 346(6210): p. 718–722.

79. Greber, B.J., et al., The complete structure of the large subunit of the mammalian mitochondrial ribosome. Nature, 2014. 515(7526): p. 283–6.

80. Itoh, Y., et al., Analysis of translating mitoribosome reveals functional characteristics of translation in mitochondria of fungi. Nat Commun, 2020. 11(1): p. 5187.

81. Rorbach, J., et al., Human mitochondrial ribosomes can switch their structural RNA composition. Proc Natl Acad Sci U S A, 2016. 113(43): p. 12198–12201.

82. Barrio-Garcia, C., et al., Architecture of the Rix1-Rea1 checkpoint machinery during pre-60S-ribosome remodeling. Nat Struct Mol Biol, 2016. 23(1): p. 37–44.

83. Datta, P.P., et al., Interaction of the G’ domain of elongation factor G and the C-terminal domain of ribosomal protein L7/L12 during translocation as revealed by cryo-EM. Mol Cell, 2005. 20(5): p. 723–31.

84. Heublein, M., et al., The novel component Kgd4 recruits the E3 subunit to the mitochondrial alpha-ketoglutarate dehydrogenase. Mol Biol Cell, 2014. 25(21): p. 3342–9.

85. Heublein, M., et al., Alternative Translation Initiation at a UUG Codon Gives Rise to Two Functional Variants of the Mitochondrial Protein Kgd4. J Mol Biol, 2019. 431(7): p. 1460–1467.

86. Obayashi, E., et al., Molecular Landscape of the Ribosome Pre-initiation Complex during mRNA Scanning: Structural Role for eIF3c and Its Control by eIF5. Cell Rep, 2017. 18(11): p. 2651–2663.

87. Ujino, S., et al., The interaction between human initiation factor eIF3 subunit c and heat-shock protein 90: a necessary factor for translation mediated by the hepatitis C virus internal ribosome entry site. Virus Res, 2012. 163(1): p. 390–5.

88. Villa, N., et al., Human eukaryotic initiation factor 4G (eIF4G) protein binds to eIF3c, -d, and -e to promote mRNA recruitment to the ribosome. J Biol Chem, 2013. 288(46): p. 32932–40.

89. Gildea, D.E., et al., The pleiotropic mouse phenotype extra-toes spotting is caused by translation initiation factor Eif3c mutations and is associated with disrupted sonic hedgehog signaling. FASEB J, 2011. 25(5): p. 1596–605.

90. Ferrari, A., S. Del’Olio, and A. Barrientos, The Diseased Mitoribosome. FEBS Lett, 2021. 595(8): p. 1025–1061.

91. Mootha, V.K., et al., Identification of a gene causing human cytochrome c oxidase deficiency by integrative genomics. Proc Natl Acad Sci U S A, 2003. 100(2): p. 605–10.

92. Xu, F., et al., The role of the LRPPRC (leucine-rich pentatricopeptide repeat cassette) gene in cytochrome oxidase assembly: mutation causes lowered levels of COX (cytochrome c oxidase) I and COX III mRNA. Biochem J, 2004. 382(Pt 1): p. 331–6.

93. Xu, F., et al., LRPPRC mutation suppresses cytochrome oxidase activity by altering mitochondrial RNA transcript stability in a mouse model. Biochem J, 2012. 441(1): p. 275–83.

94. Olahova, M., et al., LRPPRC mutations cause early-onset multisystem mitochondrial disease outside of the French-Canadian population. Brain, 2015. 138(Pt 12): p. 3503–19.

95. Fleck, D., et al., PTCD1 Is Required for Mitochondrial Oxidative-Phosphorylation: Possible Genetic Association with Alzheimer’s Disease. J Neurosci, 2019. 39(24): p. 4636–4656.

96. Rackham, O., et al., Pentatricopeptide repeat domain protein 1 lowers the levels of mitochondrial leucine tRNAs in cells. Nucleic Acids Res, 2009. 37(17): p. 5859–67.

97. Perks, K.L., et al., PTCD1 Is Required for 16S rRNA Maturation Complex Stability and Mitochondrial Ribosome Assembly. Cell Rep, 2018. 23(1): p. 127–142.

98. Newman, W.G., et al., Perrault Syndrome, in GeneReviews((R)), M.P. Adam, et al., Editors. 1993: Seattle (WA).

99. Shen, Y., et al., Geniposide Possesses the Protective Effect on Myocardial Injury by Inhibiting Oxidative Stress and Ferroptosis via Activation of the Grsf1/GPx4 Axis. Front Pharmacol, 2022. 13: p. 879870.

100. Jourdain, A.A., et al., GRSF1 regulates RNA processing in mitochondrial RNA granules. Cell Metab, 2013. 17(3): p. 399–410.

101. Giardina, G., et al., How pyridoxal 5’-phosphate differentially regulates human cytosolic and mitochondrial serine hydroxymethyltransferase oligomeric state. FEBS J, 2015. 282(7): p. 1225–41.

102. Tramonti, A., et al., Metformin Is a Pyridoxal-5’-phosphate (PLP)-Competitive Inhibitor of SHMT2. Cancers (Basel), 2021. 13(16).

103. Morscher, R.J., et al., Mitochondrial translation requires folate-dependent tRNA methylation. Nature, 2018. 554(7690): p. 128–132.

104. Colonna, M.B., et al., Functional Assessment of Homozygous ALDH18A1 Variants Reveals Alterations in Amino Acid and Antioxidant Metabolism. Hum Mol Genet, 2022.

105. Saito, Y., et al., Yeast Two-Hybrid and One-Hybrid Screenings Identify Regulators of hsp70 Gene Expression. J Cell Biochem, 2016. 117(9): p. 2109–17.

106. Dores-Silva, P.R., et al., New insights on human Hsp70-escort protein 1: Chaperone activity, interaction with liposomes, cellular localizations and HSPA’s self-assemblies remodeling. Int J Biol Macromol, 2021. 182: p. 772–784.

107. Srivastava, S., et al., Regulation of mitochondrial protein import by the nucleotide exchange factors GrpEL1 and GrpEL2 in human cells. J Biol Chem, 2017. 292(44): p. 18075–18090.

108. Muster, B., et al., Respiratory chain complexes in dynamic mitochondria display a patchy distribution in life cells. PLoS One, 2010. 5(7): p. e11910.

109. Babot, M., et al., ND3, ND1 and 39kDa subunits are more exposed in the de-active form of bovine mitochondrial complex I. Biochim Biophys Acta, 2014. 1837(6): p. 929–39.

110. Lobo-Jarne, T., et al., Multiple pathways coordinate assembly of human mitochondrial complex IV and stabilization of respiratory supercomplexes. EMBO J, 2020. 39(14): p. e103912.

111. Jackson, T.D., et al., Sideroflexin 4 is a complex I assembly factor that interacts with the MCIA complex and is required for the assembly of the ND2 module. Proc Natl Acad Sci U S A, 2022. 119(13): p. e2115566119.

112. Swenson, S., et al., Analysis of Oligomerization Properties of Heme a Synthase Provides Insights into Its Function in Eukaryotes. J Biol Chem, 2016. 291(19): p. 10411–25.

113. Taylor, N.G., et al., The Assembly Factor Pet117 Couples Heme a Synthase Activity to Cytochrome Oxidase Assembly. J Biol Chem, 2017. 292(5): p. 1815–1825.

114. International Mouse Phenotyping Consortium. see https://www.mousephenotype.org/data/genes/MGI:1858213.

115. Bareth, B., et al., The heme a synthase Cox15 associates with cytochrome c oxidase assembly intermediates during Cox1 maturation. Mol Cell Biol, 2013. 33(20): p. 4128–37.

116. Dusek, P., et al., Cerebral Iron Deposition in Neurodegeneration. Biomolecules, 2022. 12(5).

117. Wang, Z., et al., Petroleum biomarker fingerprinting for oil spill characterization and source identification. Standard Handbook Oil Spill Environmental Forensics (Second Edition), ed. W. Stout S. A. Z. Vol. Chapter 4. 2016: Academic Press.

118. Labbe, R.F., H.J. Vreman, and D.K. Stevenson, Zinc protoporphyrin: A metabolite with a mission. Clin Chem, 1999. 45(12): p. 2060–72.

119. Spasojevic, I., et al., Mn porphyrin-based superoxide dismutase (SOD) mimic, MnIIITE-2-PyP5+, targets mouse heart mitochondria. Free Radic Biol Med, 2007. 42(8): p. 1193–200.

120. Crow, J.P., et al., Manganese porphyrin given at symptom onset markedly extends survival of ALS mice. Ann Neurol, 2005. 58(2): p. 258–65.

121. Watkins, S., J. Baron, and T.R. Tephly, Identification of cobalt protoporphyrin IX formation in vivo following cobalt administration to rats. Biochem Pharmacol, 1980. 29(17): p. 2319–23.

122. Maitra, D., et al., Porphyrin-Induced Protein Oxidation and Aggregation as a Mechanism of Porphyria-Associated Cell Injury. Cell Mol Gastroenterol Hepatol, 2019. 8(4): p. 535–548.

123. Mayr, S.J., R.R. Mendel, and G. Schwarz, Molybdenum cofactor biology, evolution and deficiency. Biochim Biophys Acta Mol Cell Res, 2021. 1868(1): p. 118883.

124. Guilarte, T.R. and K.K. Gonzales, Manganese-Induced Parkinsonism Is Not Idiopathic Parkinson’s Disease: Environmental and Genetic Evidence. Toxicol Sci, 2015. 146(2): p. 204–12.

125. Vyskocil, A. and C. Viau, Assessment of molybdenum toxicity in humans. J Appl Toxicol, 1999. 19(3): p. 185–92.

126. Ohsawa, H. and C. Gualerzi, Chemical modification in situ of Escherichia coli 30 S ribosomal proteins by the site-specific reagent pyridoxal phosphate. Inactivation of the aminoacyl-tRNA and mRNA binding sites. J Biol Chem, 1983. 258(1): p. 150–6.

127. Montioli, R., et al., Deficit of human ornithine aminotransferase in gyrate atrophy: Molecular, cellular, and clinical aspects. Biochim Biophys Acta Proteins Proteom, 2021. 1869(1): p. 140555.

128. Pfanzelt, M., et al., Tailored Pyridoxal Probes Unravel Novel Cofactor-Dependent Targets and Antibiotic Hits in Critical Bacterial Pathogens. Angew Chem Int Ed Engl, 2022. 61(24): p. e202117724.

129. Montioli, R., et al., Molecular and cellular basis of ornithine delta-aminotransferase deficiency caused by the V332M mutation associated with gyrate atrophy of the choroid and retina. Biochim Biophys Acta Mol Basis Dis, 2018. 1864(11): p. 3629–3638.

130. Levchenko, I., et al., A specificity-enhancing factor for the ClpXP degradation machine. Science, 2000. 289(5488): p. 2354–6.

131. Fei, X., et al., Structural basis of ClpXP recognition and unfolding of ssrA-tagged substrates. Elife, 2020. 9.

132. Lytvynenko, I., et al., Alanine Tails Signal Proteolysis in Bacterial Ribosome-Associated Quality Control. Cell, 2019. 178(1): p. 76–90 e22.

133. Shen, B.W., et al., Crystal structure of human recombinant ornithine aminotransferase. J Mol Biol, 1998. 277(1): p. 81–102.

134. LaBreck, C.J., et al., The Protein Chaperone ClpX Targets Native and Non-native Aggregated Substrates for Remodeling, Disassembly, and Degradation with ClpP. Front Mol Biosci, 2017. 4: p. 26.

135. Truscott, K.N., A. Bezawork-Geleta, and D.A. Dougan, Unfolded protein responses in bacteria and mitochondria: a central role for the ClpXP machine. IUBMB Life, 2011. 63(11): p. 955–63.

136. Sauer, R.T. and T.A. Baker, AAA+ proteases: ATP-fueled machines of protein destruction. Annu Rev Biochem, 2011. 80: p. 587–612.

137. Snider, J. and W.A. Houry, MoxR AAA+ ATPases: a novel family of molecular chaperones? J Struct Biol, 2006. 156(1): p. 200–9.

138. Wong, K.S. and W.A. Houry, Novel structural and functional insights into the MoxR family of AAA+ ATPases. J Struct Biol, 2012. 179(2): p. 211–21.

139. Snider, J., G. Thibault, and W.A. Houry, The AAA+ superfamily of functionally diverse proteins. Genome Biol, 2008. 9(4): p. 216.

140. Felix, J., et al., The AAA+ ATPase RavA and its binding partner ViaA modulate E. coli aminoglycoside sensitivity through interaction with the inner membrane. Nat Commun, 2022. 13(1): p. 5502.

141. Wong, K.S., et al., The MoxR ATPase RavA and its cofactor ViaA interact with the NADH:ubiquinone oxidoreductase I in Escherichia coli. PLoS One, 2014. 9(1): p. e85529.

142. Wong, K.S., et al., The RavA-ViaA Chaperone-Like System Interacts with and Modulates the Activity of the Fumarate Reductase Respiratory Complex. J Mol Biol, 2017. 429(2): p. 324–344.

143. El Bakkouri, M., et al., Structure of RavA MoxR AAA+ protein reveals the design principles of a molecular cage modulating the inducible lysine decarboxylase activity. Proc Natl Acad Sci U S A, 2010. 107(52): p. 22499–504.

144. Tomar, P.C., N. Lakra, and S.N. Mishra, Cadaverine: a lysine catabolite involved in plant growth and development. Plant Signal Behav, 2013. 8(10): p. doi: 104161/psb25850.

145. Lightfoot, H.L. and J. Hall, Endogenous polyamine function--the RNA perspective. Nucleic Acids Res, 2014. 42(18): p. 11275–90.

146. Michael, A.J., Biosynthesis of polyamines and polyamine-containing molecules. Biochem J, 2016. 473(15): p. 2315–29.

147. Jessop, M., et al., Structural insights into ATP hydrolysis by the MoxR ATPase RavA and the LdcI-RavA cage-like complex. Commun Biol, 2020. 3(1): p. 46.

148. Ramirez, P., et al., Differential protein expression during growth of Acidithiobacillus ferrooxidans on ferrous iron, sulfur compounds, or metal sulfides. Appl Environ Microbiol, 2004. 70(8): p. 4491–8.

149. Tsai, Y.C., et al., Insights into the mechanism and regulation of the CbbQO-type Rubisco activase, a MoxR AAA+ ATPase. Proc Natl Acad Sci U S A, 2020. 117(1): p. 381–387.

150. Gao, Y.S., et al., Hexameric structure of the ATPase motor subunit of magnesium chelatase in chlorophyll biosynthesis. Protein Sci, 2020. 29(4): p. 1040–1046.

151. Farmer, D.A., et al., The ChlD subunit links the motor and porphyrin binding subunits of magnesium chelatase. Biochem J, 2019. 476(13): p. 1875–1887.

152. Jensen, P.E., et al., Expression of the chlI, chlD, and chlH genes from the Cyanobacterium synechocystis PCC6803 in Escherichia coli and demonstration that the three cognate proteins are required for magnesium-protoporphyrin chelatase activity. J Biol Chem, 1996. 271(28): p. 16662–7.

153. Walker, C.J. and R.D. Willows, Mechanism and regulation of Mg-chelatase. Biochem J, 1997. 327 (Pt 2)(Pt 2): p. 321–33.

154. Papenbrock, J., et al., Role of magnesium chelatase activity in the early steps of the tetrapyrrole biosynthetic pathway. Plant Physiol, 2000. 122(4): p. 1161–9.

155. Fodje, M.N., et al., Interplay between an AAA module and an integrin I domain may regulate the function of magnesium chelatase. J Mol Biol, 2001. 311(1): p. 111–22.

156. Kuznetsov, S., A. Milenkin, and I. Antonov, Translational Frameshifting in the chlD Gene Gives a Clue to the Coevolution of the Chlorophyll and Cobalamin Biosyntheses. Microorganisms, 2022. 10(6).

157. Bryant, D.A., C.N. Hunter, and M.J. Warren, Biosynthesis of the modified tetrapyrroles-the pigments of life. J Biol Chem, 2020. 295(20): p. 6888–6925.

158. Cammarano, P., et al., Characterization of unfolded and compact ribosomal subunits from plants and their relationship to those of lower and higher animals: evidence for physicochemical heterogeneity among eucaryotic ribosomes. Biochim Biophys Acta, 1972. 281(4): p. 571–96.

159. Sreedhara, A. and J.A. Cowan, Structural and catalytic roles for divalent magnesium in nucleic acid biochemistry. Biometals, 2002. 15(3): p. 211–23.

160. Yamagami, R., J.P. Sieg, and P.C. Bevilacqua, Functional Roles of Chelated Magnesium Ions in RNA Folding and Function. Biochemistry, 2021. 60(31): p. 2374–2386.

161. Welty, R., et al., Ribosomal Protein L11 Selectively Stabilizes a Tertiary Structure of the GTPase Center rRNA Domain. J Mol Biol, 2020. 432(4): p. 991–1007.

162. Akanuma, G., et al., Defect in the formation of 70S ribosomes caused by lack of ribosomal protein L34 can be suppressed by magnesium. J Bacteriol, 2014. 196(22): p. 3820–30.

163. Welty, R. and K.B. Hall, Nucleobases Undergo Dynamic Rearrangements during RNA Tertiary Folding. J Mol Biol, 2016. 428(22): p. 4490–4502.

164. Kehrein, K., et al., Organization of Mitochondrial Gene Expression in Two Distinct Ribosome-Containing Assemblies. Cell Rep, 2015. 10(6): p. 843–853.

165. Reblova, K., et al., Non-Watson-Crick basepairing and hydration in RNA motifs: molecular dynamics of 5S rRNA loop E. Biophys J, 2003. 84(6): p. 3564–82.

166. Uno, T., et al., Copper insertion facilitates water-soluble porphyrin binding to rA.rU and rA.dT base pairs in duplex RNA and RNA.DNA hybrids. Biochemistry, 2002. 41(43): p. 13059–66.

167. Wei, C., et al., Study on the interaction of porphyrin with G-quadruplex DNAs. Biophys Chem, 2008. 137(1): p. 19–23.

168. Havlova, K. and J. Fajkus, G4 Structures in Control of Replication and Transcription of rRNA Genes. Front Plant Sci, 2020. 11: p. 593692.

169. Mestre-Fos, S., et al., Profusion of G-quadruplexes on both subunits of metazoan ribosomes. PLoS One, 2019. 14(12): p. e0226177.

170. Datta, A., et al., G-Quadruplex Assembly by Ribosomal DNA: Emerging Roles in Disease Pathogenesis and Cancer Biology. Cytogenet Genome Res, 2021. 161(6-7): p. 285–296.

171. Zamiri, B., et al., TMPyP4 porphyrin distorts RNA G-quadruplex structures of the disease-associated r(GGGGCC)n repeat of the C9orf72 gene and blocks interaction of RNA-binding proteins. J Biol Chem, 2014. 289(8): p. 4653–9.

172. Weisman-Shomer, P., et al., The cationic porphyrin TMPyP4 destabilizes the tetraplex form of the fragile X syndrome expanded sequence d(CGG)n. Nucleic Acids Res, 2003. 31(14): p. 3963–70.

173. Whittaker, C.A. and R.O. Hynes, Distribution and evolution of von Willebrand/integrin A domains: widely dispersed domains with roles in cell adhesion and elsewhere. Mol Biol Cell, 2002. 13(10): p. 3369–87.

174. Romes, E.M., M. Sobhany, and R.E. Stanley, The Crystal Structure of the Ubiquitin-like Domain of Ribosome Assembly Factor Ytm1 and Characterization of Its Interaction with the AAA-ATPase Midasin. J Biol Chem, 2016. 291(2): p. 882–93.

175. Chen, Z., et al., Structural Insights into Mdn1, an Essential AAA Protein Required for Ribosome Biogenesis. Cell, 2018. 175(3): p. 822–834 e18.

176. Ulbrich, C., et al., Mechanochemical removal of ribosome biogenesis factors from nascent 60S ribosomal subunits. Cell, 2009. 138(5): p. 911–22.

177. Sano, S., et al., Significance of mitochondria for porphyrin and heme biosynthesis. Science, 1959. 129(3344): p. 275–6.

178. Luo, M., et al., Von Willebrand factor A domain-containing protein 8 (VWA8) localizes to the matrix side of the inner mitochondrial membrane. Biochem Biophys Res Commun, 2020. 521(1): p. 158–163.

179. Dietz, J.V., et al., Mitochondrial contact site and cristae organizing system (MICOS) machinery supports heme biosynthesis by enabling optimal performance of ferrochelatase. Redox Biol, 2021. 46: p. 102125.

180. Umair, M., et al., Mutated VWA8 Is Associated With Developmental Delay, Microcephaly, and Scoliosis and Plays a Novel Role in Early Development and Skeletal Morphogenesis in Zebrafish. Front Cell Dev Biol, 2021. 9: p. 736960.

181. Fujiwara, T. and H. Harigae, Biology of Heme in Mammalian Erythroid Cells and Related Disorders. Biomed Res Int, 2015. 2015: p. 278536.

182. Kim, H.J., et al., Structure, function, and assembly of heme centers in mitochondrial respiratory complexes. Biochim Biophys Acta, 2012. 1823(9): p. 1604–16.

183. Diaz, F., et al., Cytochrome c oxidase is required for the assembly/stability of respiratory complex I in mouse fibroblasts. Mol Cell Biol, 2006. 26(13): p. 4872–81.

184. Hiser, L., et al., Cox11p is required for stable formation of the Cu(B) and magnesium centers of cytochrome c oxidase. J Biol Chem, 2000. 275(1): p. 619–23.

185. Bourens, M. and A. Barrientos, A CMC1-knockout reveals translation-independent control of human mitochondrial complex IV biogenesis. EMBO Rep, 2017. 18(3): p. 477–494.

186. Timon-Gomez, A., et al., Mitochondrial cytochrome c oxidase biogenesis: Recent developments. Semin Cell Dev Biol, 2018. 76: p. 163–178.

187. Carlson, M.A., et al., Ribosomal protein L7/L12 is required for GTPase translation factors EF-G, RF3, and IF2 to bind in their GTP state to 70S ribosomes. FEBS J, 2017. 284(11): p. 1631–1643.

188. Rodnina, M.V., et al., Converting GTP hydrolysis into motion: versatile translational elongation factor G. Biol Chem, 2019. 401(1): p. 131–142.

189. Mestre-Fos, S., et al., Human ribosomal G-quadruplexes regulate heme bioavailability. J Biol Chem, 2020. 295(44): p. 14855–14865.

190. Canesin, G., et al., HO-1 and Heme: G-Quadruplex Interaction Choreograph DNA Damage Responses and Cancer Growth. Cells, 2021. 10(7).

191. Luo, M., et al., Deletion of the Mitochondrial Protein VWA8 Induces Oxidative Stress and an HNF4alpha Compensatory Response in Hepatocytes. Biochemistry, 2019. 58(49): p. 4983–4996.

192. Luo, M., et al., Deletion of Von Willebrand A Domain Containing Protein (VWA8) raises activity of mitochondrial electron transport chain complexes in hepatocytes. Biochem Biophys Rep, 2021. 26: p. 100928.

193. Greber, B.J., et al., *Ribosome.* The complete structure of the 55S mammalian mitochondrial ribosome. Science, 2015. 348(6232): p. 303–8.

194. Prezant, T.R., et al., Mitochondrial ribosomal RNA mutation associated with both antibiotic-induced and non-syndromic deafness. Nat Genet, 1993. 4(3): p. 289–94.

195. Itoh, Y., et al., Structure of the mitoribosomal small subunit with streptomycin reveals Fe-S clusters and physiological molecules. Elife, 2022. 11.

196. Amunts, A., et al., Ribosome. The structure of the human mitochondrial ribosome. Science, 2015. 348(6230): p. 95–98.

197. Jacques, C., et al., mtDNA controls expression of the Death Associated Protein 3. Exp Cell Res, 2006. 312(6): p. 737–45.

198. Hsiao, C. and L.D. Williams, A recurrent magnesium-binding motif provides a framework for the ribosomal peptidyl transferase center. Nucleic Acids Res, 2009. 37(10): p. 3134–42.

199. Sun, F.J. and G. Caetano-Anolles, The evolutionary history of the structure of 5S ribosomal RNA. J Mol Evol, 2009. 69(5): p. 430–43.

200. Gongadze, G.M., 5S rRNA and ribosome. Biochemistry (Mosc), 2011. 76(13): p. 1450–64.

201. Dontsova, O.A. and J.D. Dinman, 5S rRNA: Structure and Function from Head to Toe. Int J Biomed Sci, 2005. 1(1): p. 1–7.

202. Sloan, K.E., M.T. Bohnsack, and N.J. Watkins, The 5S RNP couples p53 homeostasis to ribosome biogenesis and nucleolar stress. Cell Rep, 2013. 5(1): p. 237–47.

203. Smirnov, A., et al., Biological significance of 5S rRNA import into human mitochondria: role of ribosomal protein MRP-L18. Genes Dev, 2011. 25(12): p. 1289–305.

204. Entelis, N.S., et al., 5 S rRNA and tRNA import into human mitochondria. Comparison of in vitro requirements. J Biol Chem, 2001. 276(49): p. 45642–53.

205. Smirnov, A., et al., Two distinct structural elements of 5S rRNA are needed for its import into human mitochondria. RNA, 2008. 14(4): p. 749–59.

206. Tobiasson, V., et al., Interconnected assembly factors regulate the biogenesis of mitoribosomal large subunit. EMBO J, 2021. 40(6): p. e106292.

207. Koper, K., et al., Evolutionary origin and functional diversification of aminotransferases. J Biol Chem, 2022. 298(8): p. 102122.

208. Liang, J., et al., Current Advances on Structure-Function Relationships of Pyridoxal 5’-Phosphate-Dependent Enzymes. Front Mol Biosci, 2019. 6: p. 4.

209. di Salvo, M.L., N. Budisa, and R. Contestabile, PLP-dependent Enzymes: a Powerful Tool for Metabolic Synthesis of Non-canonical Amino Acids. https://www.beilstein-institut.de/download/65/plp-dependent_enzymes_a_powerful_tool_for_metabolic_synthesis_of_non-canonical_amino_acids_.pdf, 2012.

210. Chen, M., C.T. Liu, and Y. Tang, Discovery and Biocatalytic Application of a PLP-Dependent Amino Acid gamma-Substitution Enzyme That Catalyzes C-C Bond Formation. J Am Chem Soc, 2020. 142(23): p. 10506–10515.

211. Ikushiro, H., et al., Heme-dependent Inactivation of 5-Aminolevulinate Synthase from Caulobacter crescentus. Sci Rep, 2018. 8(1): p. 14228.

212. Deshmukh, D.R. and S.K. Srivastava, Purification and properties of ornithine aminotransferase from rat brain. Experientia, 1984. 40(4): p. 357–9.

213. Kirschning, A., On the evolution of coenzyme biosynthesis. Nat Prod Rep, 2022. 39(11): p. 2175–2199.

214. Bottinger, L., et al., Respiratory chain supercomplexes associate with the cysteine desulfurase complex of the iron-sulfur cluster assembly machinery. Mol Biol Cell, 2018. 29(7): p. 776–785.

215. Sharma, S.S. and K.J. Dietz, The significance of amino acids and amino acid-derived molecules in plant responses and adaptation to heavy metal stress. J Exp Bot, 2006. 57(4): p. 711–26.

216. Whittaker, M.M., A. Penmatsa, and J.W. Whittaker, The Mtm1p carrier and pyridoxal 5’-phosphate cofactor trafficking in yeast mitochondria. Arch Biochem Biophys, 2015. 568: p. 64–70.

217. Smith, T.F. and H. Hartman, The evolution of Class II Aminoacyl-tRNA synthetases and the first code. FEBS Lett, 2015. 589(23): p. 3499–507.

218. Schneider, A., T. Stachelhaus, and M.A. Marahiel, Targeted alteration of the substrate specificity of peptide synthetases by rational module swapping. Mol Gen Genet, 1998. 257(3): p. 308–18.

219. Mills, P.B., et al., Epilepsy due to PNPO mutations: genotype, environment and treatment affect presentation and outcome. Brain, 2014. 137(Pt 5): p. 1350–60.

220. Henslee, J.G., et al., Factors influencing pyrroline 5-carboxylate synthesis from glutamate by rat intestinal mucosa mitochondria. Arch Biochem Biophys, 1983. 226(2): p. 693–703.

221. Csonka, L.N. and T. Leisinger, Biosynthesis of Proline. EcoSal Plus, 2007. 2(2).

222. Zappa, S., K. Li, and C.E. Bauer, The tetrapyrrole biosynthetic pathway and its regulation in Rhodobacter capsulatus. Adv Exp Med Biol, 2010. 675: p. 229–50.

223. Hoober, J.K., et al., Biosynthesis of delta-aminolevulinate in greening barley leaves. IX. Structure of the substrate, mode of gabaculine inhibition, and the catalytic mechanism of glutamate 1-semialdehyde aminotransferase. Carlsberg Res Commun, 1988. 53(1): p. 11–25.

224. Sinha, N., et al., Glutamate 1-semialdehyde aminotransferase is connected to GluTR by GluTR-binding protein and contributes to the rate-limiting step of 5-aminolevulinic acid synthesis. Plant Cell, 2022. 34(11): p. 4623–4640.

225. Zhang, X., et al., Translational control of the cytosolic stress response by mitochondrial ribosomal protein L18. Nat Struct Mol Biol, 2015. 22(5): p. 404–10.

226. Kim, H.K., S.I. Choi, and B.L. Seong, 5S rRNA-assisted DnaK refolding. Biochem Biophys Res Commun, 2010. 391(2): p. 1177–81.

227. Brusslan, J.A. and M.P. Peterson, Tetrapyrrole regulation of nuclear gene expression. Photosynth Res, 2002. 71(3): p. 185–94.

228. Barros, M.H., et al., Involvement of mitochondrial ferredoxin and Cox15p in hydroxylation of heme O. FEBS Lett, 2001. 492(1-2): p. 133–8.

229. Antonicka, H., et al., Mutations in COX15 produce a defect in the mitochondrial heme biosynthetic pathway, causing early-onset fatal hypertrophic cardiomyopathy. Am J Hum Genet, 2003. 72(1): p. 101–14.

230. Antonicka, H., et al., Mutations in COX10 result in a defect in mitochondrial heme A biosynthesis and account for multiple, early-onset clinical phenotypes associated with isolated COX deficiency. Hum Mol Genet, 2003. 12(20): p. 2693–702.

231. Alfadhel, M., et al., Infantile cardioencephalopathy due to a COX15 gene defect: report and review. Am J Med Genet A, 2011. 155A(4): p. 840–4.

232. Zheng, H., et al., Molecular cloning and characterization of a novel human putative transmembrane protein homologous to mouse sideroflexin associated with sideroblastic anemia. DNA Seq, 2003. 14(5): p. 369–73.

233. Paul, B.T., et al., Sideroflexin 4 affects Fe-S cluster biogenesis, iron metabolism, mitochondrial respiration and heme biosynthetic enzymes. Sci Rep, 2019. 9(1): p. 19634.

234. Hildick-Smith, G.J., et al., Macrocytic anemia and mitochondriopathy resulting from a defect in sideroflexin 4. Am J Hum Genet, 2013. 93(5): p. 906–14.

