## Supplementary figures and images for "Translation fidelity and respiration deficits in CLPP-deficient tissues: Mechanistic insights from mitochondrial complexome"

### FigureS1-VWA8human-InteractingPreysBIOGRID.tif

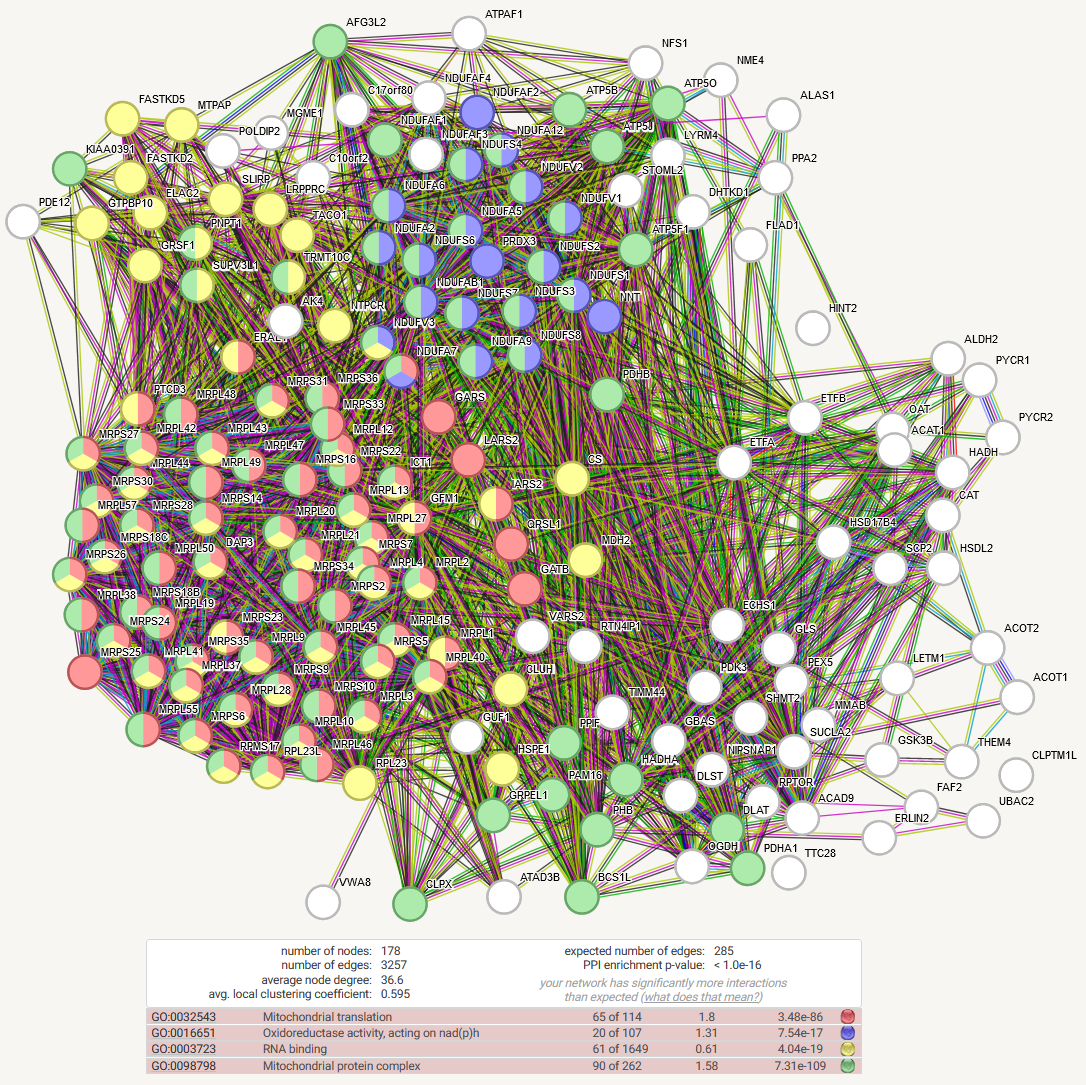

### FigureS2-CLPX-VWA8-Ribonucleoproteins.tif

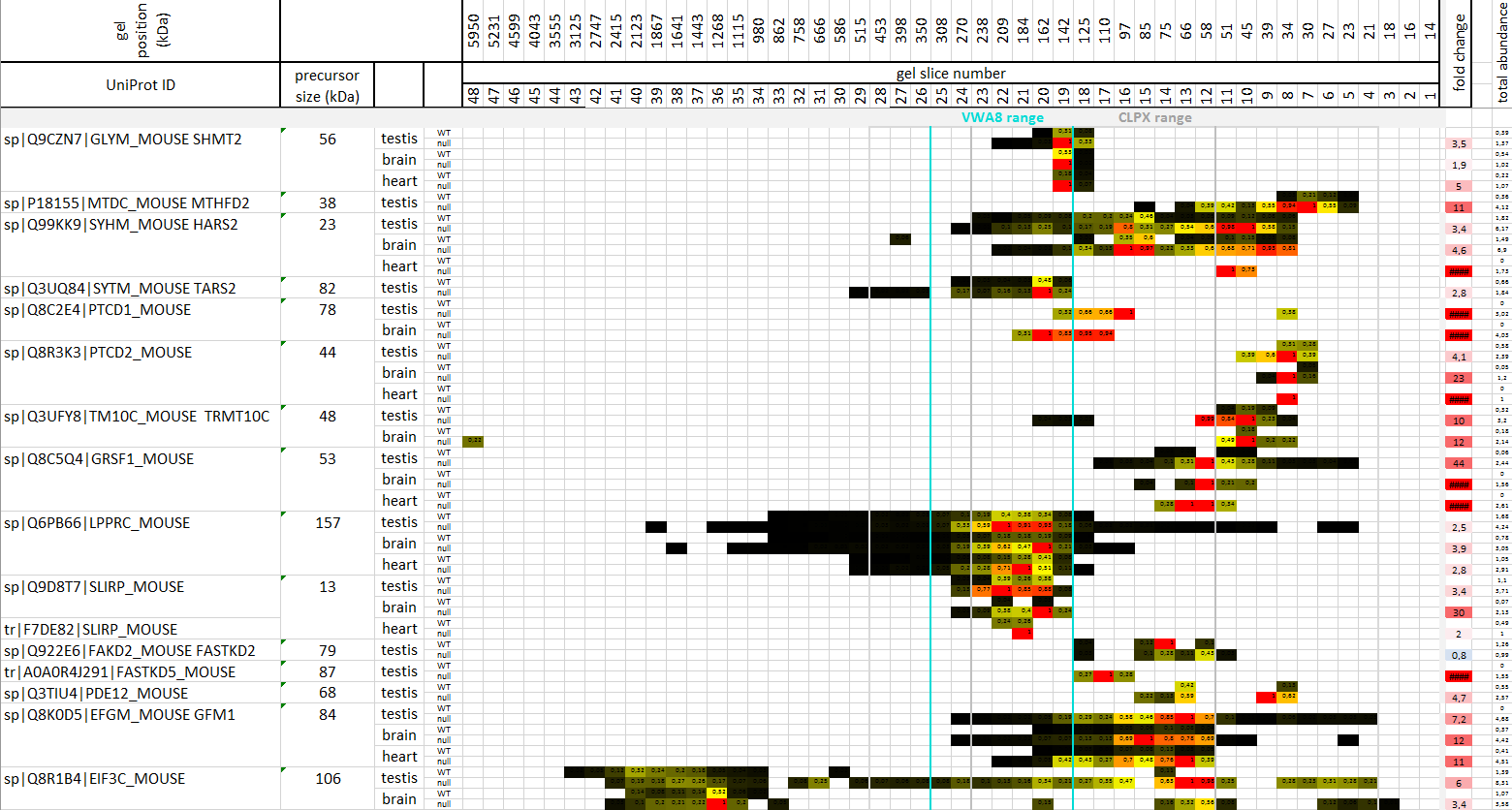

### FigureS3-CLPX-VWA8-ComigratingDisperseAccumulated-PLPfactors.tif

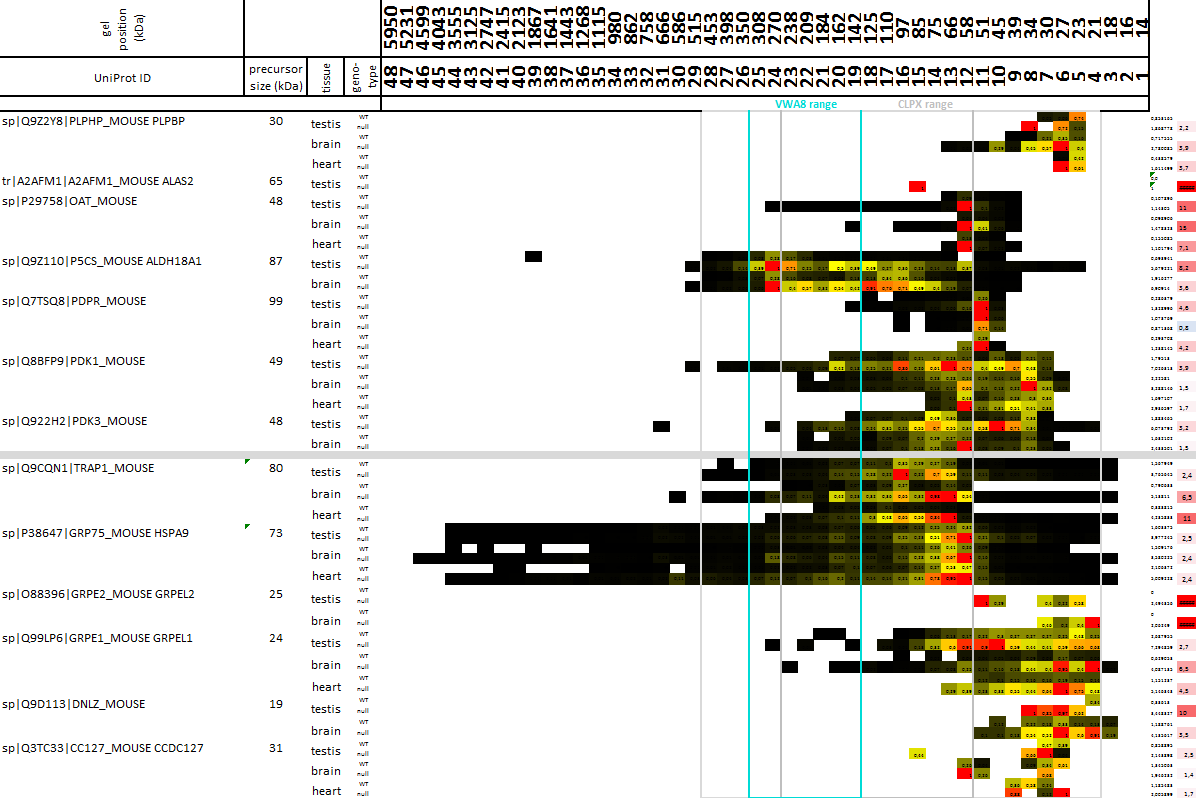

### FigureS4-CLPX-VWA8-ComigratingDisperseAccumulated.tif

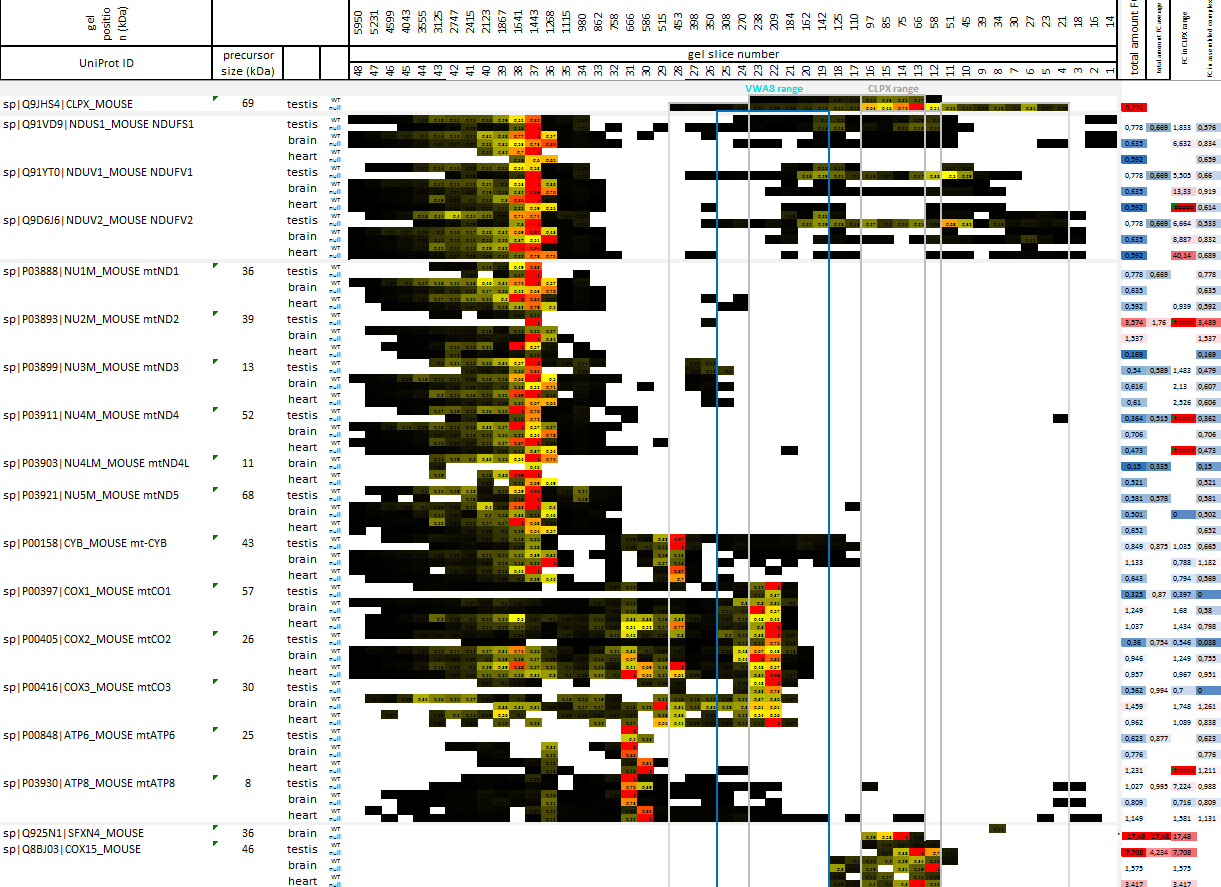
